# Signal Integration and Adaptive Sensory Diversity Tuning in *Escherichia coli* Chemotaxis

**DOI:** 10.1101/2023.02.08.527720

**Authors:** Jeremy Philippe Moore, Keita Kamino, Rafaela Kottou, Thomas Shimizu, Thierry Emonet

## Abstract

In uncertain environments, phenotypic diversity can be advantageous for survival. However, as the environmental uncertainty decreases, the relative advantage of having diverse phenotypes decreases. Here, we show how populations of E. coli integrate multiple chemical signals to adjust sensory diversity in response to changes in the prevalence of each ligand in the environment. Measuring kinase activity in single cells, we quantified the sensitivity distribution to various chemoattractants in different mixtures of background stimuli. We found that when ligands bind uncompetitively, the population tunes sensory diversity to each signal independently, decreasing diversity when the signal ambient concentration increases. However, amongst competitive ligands the population can only decrease sensory diversity one ligand at a time. Mathematical modeling suggests that sensory diversity tuning benefits *E. coli* populations by modulating how many cells are committed to tracking each signal proportionally as their prevalence changes.

## Introduction

While navigating their environments, organisms sense and respond to signals embedded in a complex backdrop of other stimuli. Behavioral decisions thus require sensory systems capable of parsing these rich signal mixtures to control subsequent responses. In fly olfaction for example, relatively few olfactory receptors (∼50) are used to encode and identify an enormous range of stimuli to drive a highly olfaction-dependent lifestyle.^1-4^ Similarly, many bacteria use a small set (∼3) of quorum-sensing receptors to drive the switch between biofilm and planktonic lifestyles as the ratio of signals secreted by self and other bacteria changes.^5,6^ In both of these cases, a downstream network – neural for the fly and transcriptional for quorum sensing – parses the integrated signals and prevents signal cross-talk.

One of the most-studied sensory systems for navigation is the chemotaxis network of *Escherichia coli*. In this relatively simple signaling network, five chemoreceptor species that mix within allosterically coupled signaling complexes detect many different extracellular chemicals.^7^ Ligand-receptor binding then modulates the activity of the kinase CheA, the sole output of the complex, which phosphorylates a response regulator CheY that controls the direction of flagellar rotation.^7,8^ Thus, a multitude of extracellular signals are integrated into the activity of a single kinase and downstream effectors. In these receptor-kinase complexes (Figure 1A), ligand binding at one receptor is thought to not only change the conformation of the kinase associated with that receptor, but also its neighbors, leading to amplification of the signal generated by small changes in ligand concentration.^9,10^

**Figure 1.**
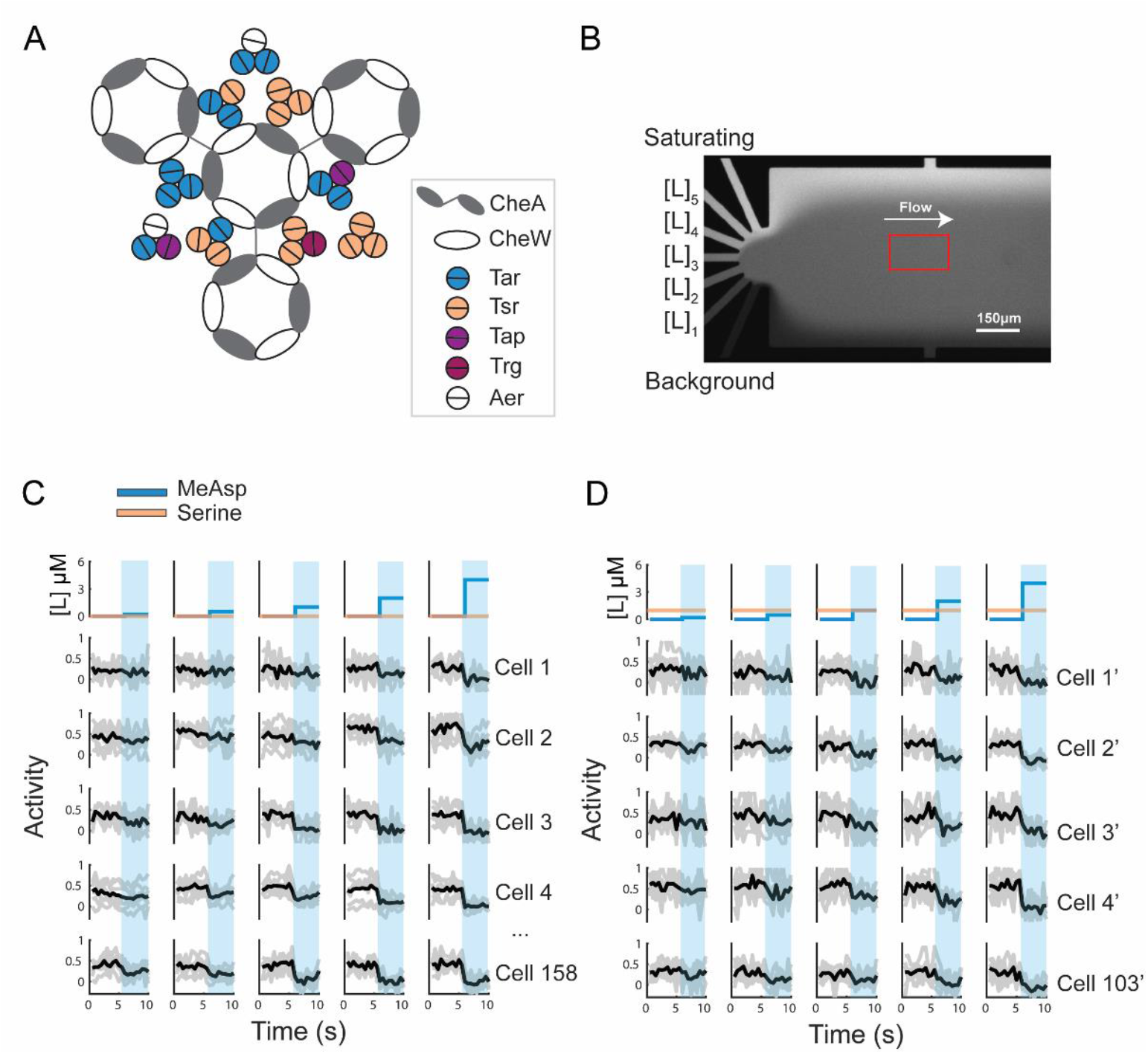
Measuring chemosensory response variation with various combinations of background and foreground ligands. **A)** Diagram of the *E. coli* chemoreceptor cluster. *E. coli* have five chemoreceptor species, but the aspartate (Tar) and serine (Tsr) receptors make up the majority of the cluster. All receptors modulate activity of the kinase CheA, which is arranged in a hexagonal array, with adjacent kinases linked by the coupling protein CheW. **B)** Single-cell FRET Measurements were performed in a microfluidic device capable of exchanging media in the field-of-view on the order of 200 ms. The device has seven inlets: one contains only the background stimulus, one a saturating stimulus to determine maximal response amplitude, and five different test stimulus levels. Flow was visualized by loading different concentrations of fluorescein in each channel. **C)** Single cell responses to five different MeAsp concentrations with no background ligands present. For each stimulus level (columns), seven responses were measured per cell. Individual responses are in gray, with the within-cell averages shown in black. Responses are normalized to the minimum and maximum FRET response measured at the beginning and end of each experiment. **D)** Example responses to MeAsp after adaptation to a 1 µM serine background.

In addition to cooperative receptor clusters, the chemotaxis network includes two proteins (CheR and CheB) that post-translationally modify receptors by respectively adding and removing methyl groups, to increase or decrease their activity in a way that opposes the current state of CheA activity.^7^ This negative feedback allows kinase activity to precisely adapt to sustained changes in attractant concentration, returning to the same steady-state activity over a wide concentration range.^7,11,12^ Adaptation has been shown to only be precise in receptor clusters containing multiple receptor species.^13^ Additionally, transient cross-methylation of receptors in the presence of a non-cognate ligand can occur.^14-16^ Despite decades of study, our understanding of cooperativity, adaptation, and signal integration within chemosensory arrays comprising multiple receptor species remains far from complete. Consequently, it remains challenging to explain, let alone predict, how *E. coli* navigate complex environments where they encounter and adapt to many signals at the same time.

Isogenic cell populations could generate individuals with diverse sensitivity to different stimuli to hedge their bets against changes in available attractant gradients and effectively explore complex environments.^17-19^ Developments in single-cell fluorescence resonance energy transfer (FRET) have allowed this diversity to be probed at the signal-transduction level by measuring FRET between CheY-mRFP1 and CheZ-YFP fusions to quantify kinase activity in individual cells.^20-23^ These studies revealed diverse sensitivities to various attractants, and confirmed cell-to-cell variability in kinase activity fluctuation.^24-26^

Recently, using single-cell FRET to measure kinase activity, we discovered that the degree of cell-to-cell variation in sensitivity to an attractant strongly depends on its background concentration.^27^ When the background concentration of a chemoattractant is low, sensitivity to the attractant varies greatly from cell to cell. However, after the cells have adapted to a sufficiently high background of the attractant, the degree of variation in sensitivity within the clonal population is strongly attenuated. This background stimulus dependent “diversity tuning” occurs without changes in gene expression. It was hypothesized that diversity tuning could play a role in a population’s ability to switch between different navigational strategies depending on the availability of environmental cues. When the information about the environment is scarce, populations can hedge their bets by diversifying their sensitivities to stimuli. However, after detecting a sufficiently strong signal, the population can shift to a tracking strategy where each individual is tuned to the ambient signal level, and primed to detect changes from the background signal level.

The diversity-tuning phenomenon was observed for the responses to alpha-methyl-aspartate (MeAsp) and serine, cognate ligands of the two major chemoreceptors Tar and Tsr, respectively, in settings where only a single ligand species is presented to cells at a time. However, whether and how diversity tuning applies to more ecologically relevant scenarios, where multiple ligands are present simultaneously, remains unknown. Does adding one stimulus to the environment affect the ability of the population to modulate the sensitivity distribution for other stimuli? Are there different effects when the ligands presented bind the same receptor or different receptors?

We hypothesized that, due to strong cooperative interactions within the receptor cluster^28^, the presence of one background ligand species could affect diversity-tuning of the sensitivity to other (foreground) ligand species. To test this hypothesis, we combined microfluidics and single-cell FRET to measure the distribution of sensitivities for various foreground ligands after allowing cells to adapt to different background conditions. We further reasoned that the binding mode of the foreground and background ligands – binding different receptors, binding one receptor competitively, or binding one receptor uncompetitively – could play an important role. Accordingly, we performed experiments for all three cases.

We found that the sensitivity distributions for ligands which bind uncompetitively, whether on the same receptor or different, despite strong allosteric coupling in the receptor cluster, can be tuned independently. However, when ligands compete for binding sites, response diversity that has collapsed upon adaptation with respect to one ligand can be restored by addition of a competitive ligand to the adapted-state background. Lastly, we explore through mathematical modeling the consequences of diversity tuning on chemotactic performance during navigation and how similar diversity tuning might arise in other systems.

## Results

### Measuring the distribution of chemotactic response sensitivities

We set out to measure the distribution of sensitivity for one ligand species in the presence of different background ligand species. In our experimental setup, the ligand whose concentration is dynamically modulated in an experiment is called the foreground ligand, and the ligand whose concentration is fixed to a constant level is called the background ligand. Following preceding works,^21,27^ we defined ‘kinase activity’, *a*, as the relative FRET level normalized within individual cells: zero and one activity correspond to the minimum and maximum FRET level, respectively. The minimum FRET levels are observed immediately following addition of a stimulus large enough to saturate the response, and the maximum FRET is observed after removal of this stimulus. We quantified sensitivity for a foreground ligand by the *K*_1/2_, which we defined as the ligand concentration where kinase activity reached half of its steady-state value. For simplicity, we focused on ligands that bind the two highest-expressed receptors in *E. coli*: Tar and Tsr.^7^

For a given combination of foreground and background stimuli, we measured the *K*_1/2_ distribution using an extension of previous methods, where cells expressing sticky FliC (FliC*) adhered to the bottom of a microfluidic device are exposed to constant flow of stimuli dissolved in motility buffer (STAR Methods).^27^ We developed a microfluidic device with seven input channels (Figure 1B; Figure S1A-D) which allowed us to alternate between flows of six different stimulus solutions and a background solution over a field of immobilized cells. Each stimulus solution was presented to cells multiple times in a single experiment to reduce the statistical uncertainties originating from measurement noise and temporal variations in kinase activity by allowing within-cell averaging of the FRET responses to each stimulus (Figure 1C, D). The time to fully exchange the media in a field of view is ∼200 ms (Figure S1A-D), enabling quantification of kinase activity responses to small stimuli without the confounding effects of response adaptation, which occurs on the order of 10 seconds for sub-saturating step stimuli.^22^ Each foreground stimulus lasted for 5 seconds with time-lapse FRET measurements conducted every 0.5 seconds starting 5 seconds before the stimulus onset and lasting 10 seconds total. We waited 30 seconds between consecutive foreground stimuli to allow kinase activity to adapt back to its steady-state level. Cells were not imaged during these adaptation periods to minimize photobleaching. This stimulus protocol gives quasi-stationary distributions of responses over the course of an experiment (Figure S2A). Data collected from at least two biological replicates were included in each analysis to increase the number of cells per data point.

Using this experimental setup, we measured the *K*_1/2_ cumulative distribution function (CDF) for one combination of foreground and background ligands in a single experiment. The CDF of *K*_1/2_ can be extracted from the distribution of post-stimulus kinase activity normalized by its steady-state level, *R* (Figure S1F). As described previously, for a population of bacteria, the value of the CDF of the *K*_1/2_ is given by the fraction of responses *R* smaller than 0.5 (Figure S1E, F).^27^

Also in agreement with previous work, the *K*_1/2_ CDFs were well approximated by a lognormal distribution (Figure S2B, C).^27^ To quantify the variation in the magnitude of sensory diversity we use the coefficient of variation (CV) of the *K*_1/2_ distribution, which is defined as the standard deviation divided by the mean. Note that the 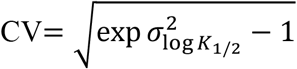 is also monotonically related to the standard deviation 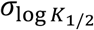 of the distribution of log *K*_1/2_.

### Orthogonal diversity tuning for ligands that bind different receptors

We first investigated how adaptation to a background ligand that binds one receptor species affects the *K*_1/2_ distribution for foreground ligands that bind a different receptor species (Figure 2A). Previous population-averaged FRET experiments found that prior adaptation to serine, which binds Tsr with high affinity (*K*_1/2_ ≈ 10^−1^µM) and Tar with low affinity (*K*_1/2_ ≈ 10^1^µM), does not significantly affect the population-averaged *K*_1/2_ of MeAsp, which primarily binds Tar.^29,30^ However, at the single-cell level, whether this independence extends not only to the mean but also the variance of the *K*_1/2_ distribution remains unknown.

**Figure 2.**
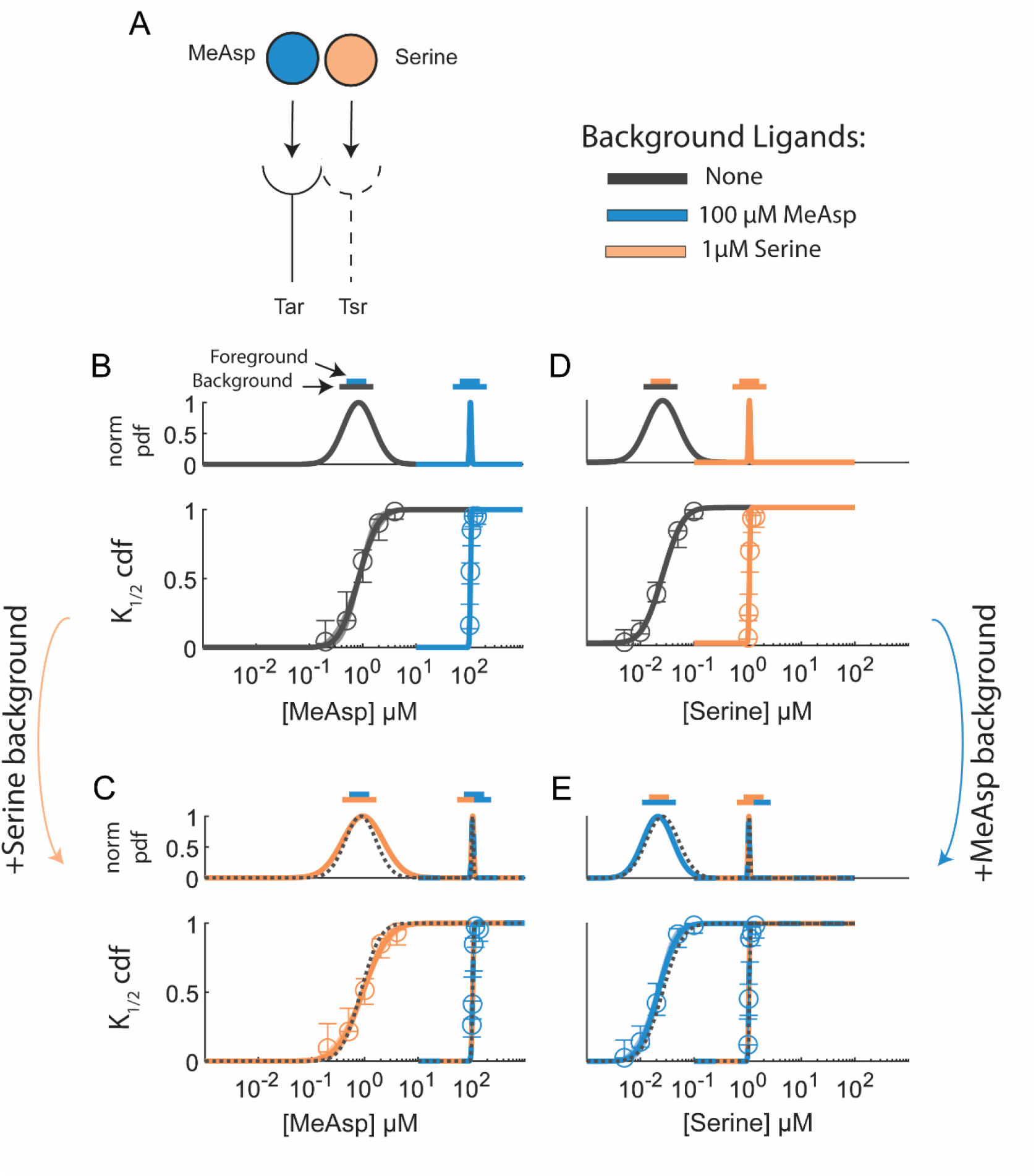
Responses to ligands that bind different receptors show independent diversity tuning. **A)** We measured responses to MeAsp and serine while varying the background concentration of both ligands. **B)** Cumulative distribution function (cdf) of the MeAsp *K*_1/2_ measured in plain buffer (gray), and after adaptation to a 100 *µM* MeAsp background stimulus (blue). Points on the cdf are determined by calculating the fraction of cells with a response less than half the steady-state activity at each concentration (*N* = 153, 171, respectively for each curve as they appear from left to right, where *N* is the number of cells used to confine each cdf. All sample sizes will reported as such unless otherwise stated). Open circles and error bars are the median and 95% confidence interval (CI) estimates from bootstrapping over cells. Parameters of the lognormal cdf and their confidence intervals are determined using a Bayesian method with uninformative priors (see Figure S2). Curves are the cdf with maximum a posteriori parameters, with shading representing the 95% CI of the cdf. The probability density function (pdf), normalized by its maximum value (norm pdf) is included as a visual guide, with double-bar cartoons above each pdf to indicate the foreground ligand (top bar) and background condition (bottom bar). In all subsequent figures, *K*_1/2_ distributions should be interpreted as described above. **C)** MeAsp *K*_1/2_ distributions after adaptation to 1 *µM* serine. Background serine had no significant effect on the MeAsp *K*_1/2_ distribution with or without 100 µM MeAsp in the background (*N* = 108, 46; see Figure S2 for CV confidence intervals). Dotted line is the curve from panel B reproduced for comparison. **D)** Serine *K*_1/2_ distribution before and after adaptation to a 1 µM serine background (*N* = 174, 103). **E)** Serine *K*_1/2_ distributions after adaptation to 100 µM MeAsp (*N* = 79, 75). Black dotted lines are curves from panel D, reproduced for comparison.

To address this question, we measured the *K*_1/2_ distribution for MeAsp and serine in four background conditions: no-background (plain motility buffer), 100 µM MeAsp, 1 µM serine, and a mixture of 100 µM MeAsp and 1 µM serine. These concentrations were chosen because 100 µM MeAsp and 1 µM serine were previously shown to collapse the width of MeAsp and serine *K*_1/2_ distributions respectively upon adaptation to a background of the same signals.^27^

We observed a broad *K*_1/2_ distribution at zero-background (CV = 0.74 ± 0.12), and adaptation to 100 µM MeAsp led to a collapse in the *K*_1/2_ diversity (CV = 0.037 ± 0.004) for MeAsp (Figures 2B; Figure S2DE). However, adaptation to 1 µM serine did not significantly affect the *K*_1/2_ distribution for MeAsp, even in the mixed background condition (Figure 2C; Figure S2FG). Similarly, while adaptation to 1 µM serine background collapses the serine *K*_1/2_ diversity (Figures 2D), adaptation to 100 µM MeAsp did not significantly affect the serine *K*_1/2_ distributions (Figures 2E; Figure S2FG).

These experiments show that in addition to maintaining the same mean sensitivity for MeAsp (serine) in the presence of serine (MeAsp), the bacterial populations also maintain the degree of variability in sensitivity. As such, even though receptor clusters inside individual cells comprise a mixture of receptor species that signal cooperatively, populations of *E. coli* can independently tune the *K*_1/2_ distributions for ligands that bind different receptor species.^10,16,31^

### Diversity tuning when ligands bind the same receptor

Having seen that two ligands that primarily bind different receptor species have no effect on each other’s *K*_1/2_ distributions, we next asked how *K*_1/2_ diversity is affected when background and foreground ligands bind the same receptor. To that end, we sought to measure the *K*_1/2_ distribution for MeAsp before and after adaptation to other Tar-binding ligands.

The amino acids L-aspartate (L-asp), glutamate, and MeAsp all bind directly and competitively to the Tar receptor (Figure 3A). We first measured the *K*_1/2_ distribution for each of these ligands with no background stimulus, and found that, despite differences in the average sensitivity for each ligand, the degree of *K*_1/2_ diversity was similar for each ligand (Figure 3B, see Figure S3A-C for plots of the CV values). We then measured the MeAsp *K*_1/2_ distribution after adaptation to various concentrations of glutamate or L-asp (Figure 3C; Figure S3D-F). Although the average *K*_1/2_ for MeAsp increased in the presence of these competing background ligands, there was no significant change in the MeAsp *K*_1/2_ CV (Figure S3E). While adaptation to an L-asp or glutamate background will lead to methylation of the Tar receptor, these receptor modifications appear to have had no effect on *K*_1/2_ diversity for MeAsp.

**Figure 3.**
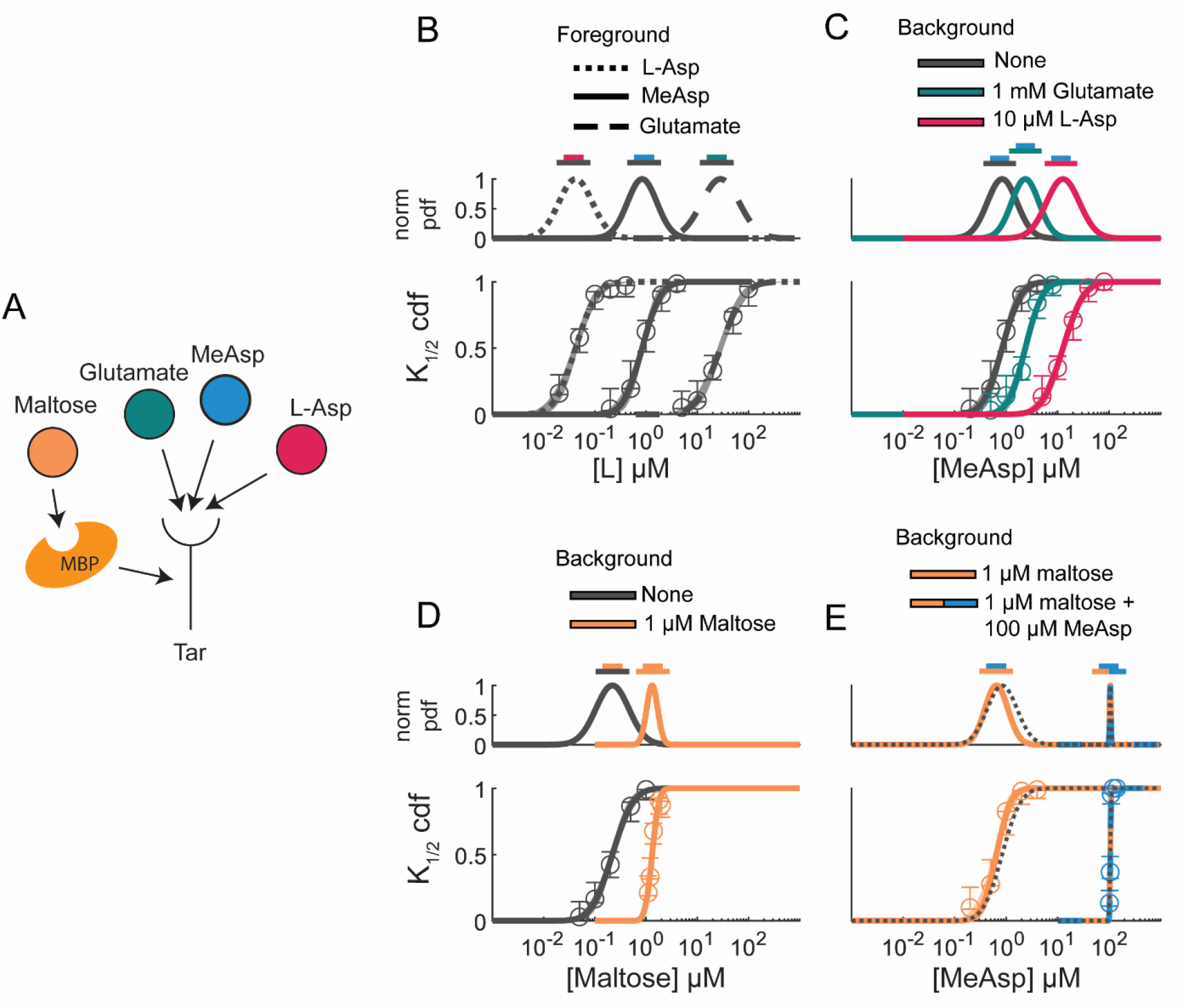
Diversity tuning when multiple ligands bind the same receptor. **A)** We measured the *K*_1/2_ distribution for 3 ligands that compete with MeAsp at the Tar ligand binding pocket, and maltose which binds Tar indirectly through the maltose binding protein (MBP). **B)** *K*_1/2_ distributions for L-Asp, MeAsp and glutamate (*N* = 108, 158, 117) measured in the absence of a background stimulus. **C)** The MeAsp *K*_1/2_ distribution in the presence of glutamate or L-asp (*N* = 158, 127, 115). In the presence of competitive background ligands, the average *K*_1/2_ increases, but the width of the curves does not collapse. There is no significant change in the CV (see Figure S3). **D)** *K*_1/2_ distribution for maltose in buffer, and after adaptation to 1 *µM* maltose (*N* = 104, 134). Even though it binds indirectly, the maltose *K*_1/2_ distribution is still tuned by adaptation to maltose. **E)** At 1 µM, Maltose does not significantly affect the mean or CV of the MeAsp *K*_1/2_ distribution (*N* = 102, 136; see Figure S3). The black dotted lines are reproduced from Figure 2B for comparison.

To separate the effects of receptor modification and competition for binding sites, we measured responses to MeAsp after adaptation to maltose, which binds Tar indirectly through the maltose binding protein. Population-averaged maltose responses are known to be independent of the ambient MeAsp concentration (Figure 3D).^29^ We first verified that responses to maltose undergo diversity tuning similar to directly binding ligands. In the absence of background stimuli, variation in maltose *K*_1/2_ was similar to that of MeAsp (CV = 0.85 ± 0.11; Figure 3E; Figure S3A-C). After adaptation to 1 µM maltose, there was a significant decrease in the Maltose *K*_1/2_ CV (CV = 0.25 ± 0.026; Figure S3B). We attempted to measure *K*_1/2_ distributions at higher backgrounds, but due to saturation of the maltose binding protein, were unable to detect responses above 10 µM, consistent with previous population-level assays.^29^

When we measured the MeAsp *K*_1/2_ distribution in the presence of 1 µM maltose, we found no significant change in the MeAsp *K*_1/2_ distribution (CV = 0.55 ± 0.12; Figure 3E; Figure S3E). Similar to when serine, which binds at a different receptor, was present in the background, even in a mixed 100 µM MeAsp + 1 µM maltose background, the MeAsp *K*_1/2_ distribution was similar to when there was a 100 µM MeAsp background alone (CV = 0.026 ± 0.002; Figure 3E). These results suggest that the shift in average *K*_1/2_ when the background is competitive is due to lower binding site availability and not receptor modification (see model and discussion below).

Taken together, our experiments with Tar-binding background ligands suggest that adaptation to a background ligand that binds one receptor, only affects the *K*_1/2_ distribution of other ligands which bind the same receptor through competition for binding sites. This is in contrast to the case where the ambient concentration of the foreground ligand itself is increasing, which leads to a decrease in *K*_1/2_ diversity. To interpret this result, we turned to mathematical modeling of the chemoreceptor cluster.

### The standard allosteric model of chemotaxis captures *K*_1/2_ diversity tuning with mixed background stimuli

We asked to what extent our results can be recapitulated by the simple Monod-Wyman-Changeux (MWC) modeling framework for chemoreceptor clusters, which has been the standard model of chemotaxis for the past decade.^32-35^ In this model, receptor-kinase clusters switch in an all-or-none fashion between a kinase activating ‘on’ state, and a deactivating ‘off’ state (**Figure 4A**). The free energy difference between these two states is determined by the concentration of all ligands in the environment, the methylation level of the receptors, and the degree of cooperativity between neighboring receptor-kinase units. Stronger cooperativity between neighboring receptor-kinase units leads to larger gain and a more nonlinear response.^9,10,36^ The model assumes precise adaptation of kinase activity,^37,38^ i.e., the change in free energy due to methylation always compensates for prolonged changes in ambient ligand concentration, ensuring that the average steady-state kinase activity is independent of the background ligand concentration. We also assume that the cluster contains two receptor species Tar and Tsr, that contribute *n*_*tar*_ and *n*_*tsr*_ cooperative units to the cluster.

**Figure 4.**
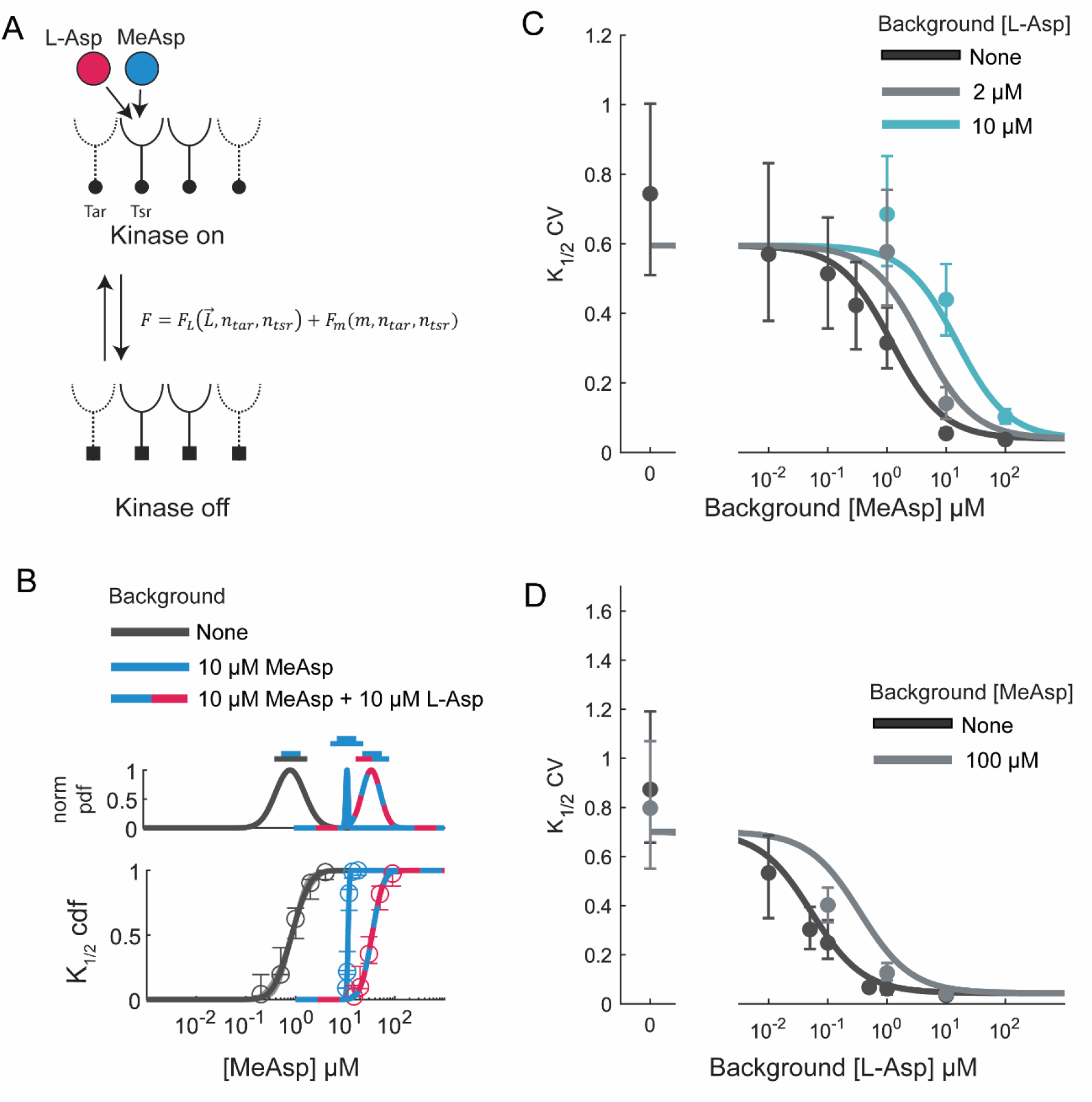
Diversity tuning in mixed backgrounds is captured by a simple allosteric model. **A)** The Monod-Wyman-Changeux (MWC) model of allostery treats receptor clusters as a single unit with ‘kinase on’ and ‘kinase off’ states. The equilibrium activity is determined by the total free energy of the cluster *F* which is decomposed into a term *F*_*L*_(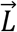, *n*_*Tar*_, *n*_*Tsr*_) which accounts for changes in the free energy due to ligand binding, and *F*_*m*_(*m, n*_*tar*_, *n*_*tsr*_) which accounts for changes in free energy due to receptor methylation. We assume precise adaptation, where at steady-state, changes in *F*_*L*_(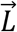, *n*_*Tar*_, *n*_*Tsr*_) are perfectly offset by *F*_*m*_(*m, n*_*tar*_, *n*_*tsr*_). **B)** MeAsp *K*_1/2_ distribution in three background conditions: Zero background (reproduced from Figure 2B for comparison), 10 *µM* MeAsp, and a mixture of 10 *µM* MeAsp with 10 *µM* L-Asp (*N* = 158, 102, 83). With background MeAsp alone, the *K*_1/2_ distribution collapses. The addition of L-Asp to this background expands the *K*_1/2_ distribution. Equation (1) was fit simultaneously to the **C)** MeAsp *K*_1/2_ CV and **D)** L-asp *K*_1/2_ CV measured in various mixed MeAsp and L-asp backgrounds. Solid points are maximum a-posteriori estimates of the CV, with error bars representing 95% confidence intervals. In both plots, the x axis represents the background concentration of MeAsp or L-asp respectively, with the color indicating the background concentration of L-asp or MeAsp respectively. The model was fit with three free parameters (mean zero-background *K*_1/2_, standard deviation in zero-background *K*_1/2_, and receptor-ligand affinity) for each ligand species, for a total of six parameters (See Table S1).

In a previous study, we showed that when just one ligand species is used as both the foreground and background stimulus, the MWC model can quantitatively recapitulate the observed *K*_1/2_ distributions of adapted populations by assuming cell-to-cell variation in only one parameter, the number of cooperative units of the cognate receptor (*n*_*tar*_ for MeAsp, *n*_*tsr*_ for serine).^27^ Here, we ask what this model predicts for the *K*_1/2_ CV in the more general case where the background can be a mixture of multiple ligand species. Motivated by our measurements, we solved the MWC model for the CV of the *K*_1/2_ distribution for a foreground ligand *L*, as a function of its background concentration *L*_0_ as well as the concentrations of all other competitive ligands *L*_*l*_ (with *l* = 1, 2, 3, …) present in the background (Figure 4B, STAR Methods):

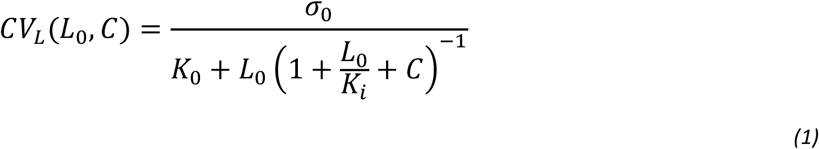

where *σ*_0_ and *K*_0_ are the standard deviation and mean of the *K*_1/2_ distribution for the foreground ligand in the absence of any background stimulus, *K*_*i*_ is the dissociation constant of the foreground ligand for the inactive receptor state, and *C* = ∑_*l*_ *L*_*l*_/*K*_*l*_ is the sum of all competing background ligand concentrations scaled by their respective dissociation constants *K*_*l*_ (STAR Methods).

This simple expression for response diversity derived from the standard MWC model of bacterial chemotaxis captures the essential features of our experiments.^33^ First, the CV of the *K*_1/2_ distribution does not depend on the concentration of uncompetitive ligands because their effect is canceled out by adaptation (STAR Methods). The concentrations *L*_*l*_ of uncompetitive ligands therefore do not appear in equation (1), which implies that they should have no effect on each other’s *K*_1/2_ diversity, as experimentally demonstrated in Figure 2CE. Second, equation (1) predicts that when the foreground ligand species is absent from the background (*L*_0_ = 0), the CV and hence *K*_1/2_ diversity should be invariant under changes in either the dissociation constant *K*_*i*_ or the amount of competitive background ligands *C*, consistent with our experiments in Figure 3BC.

Equation (1) makes several predictions. For a population adapted to a background ligand concentration *L*_0_ sufficient to collapse the *K*_1/2_ diversity for the ligand *L*, addition of a competitor to the background (positive change in C) should actually increase *K*_1/2_ diversity. To test this prediction, we measured the MeAsp *K*_1/2_ distribution after adaptation to two different backgrounds: 10 μM MeAsp alone, and 10 μM MeAsp in combination with 10 μM L-Asp (Figure 4B). Adapting to 10 μM MeAsp alone led to a collapse of the *K*_1/2_ diversity (CV = 0.055 ± 0.007). However, in the mixed background, the *K*_1/2_ distribution widened (CV = 0.44 ± 0.05), qualitatively consistent with the model.

To test the quantitative agreement between model and data, we measured the MeAsp *K*_1/2_ CV and L-asp *K*_1/2_ CV across a range of mixed backgrounds containing different concentrations of MeAsp and L-asp. We jointly fit the model to both datasets (Figure 4CD) with the *K*_0_, *σ*_0_, and *K*_*i*_ values for MeAsp and L-asp (6 parameters total) as free parameters. We generally found good agreement between the data and the model fit to both datasets (Figure 4CD) with reasonable parameter values (Table S1). Using the model, we were able to predict the full *K*_1/2_ distributions for MeAsp and L-asp in most background conditions (Figure S4). The model slightly underpredicts the mean *K*_1/2_ for MeAsp in high L-asp concentrations (Figure S4J-L), and the diversity of L-asp *K*_1/2_ values with MeAsp present (Figure 4D). *E. coli* were previously shown to respond to L-asp even with mutant Tar receptors lacking the ligand binding domain,^30^ possibly through interactions with the phosphotransferase system.^39^ Such effects are not accounted for by the MWC model, and may be a source of the deviations from the model we observe.

### Impact of sensory diversity tuning on navigation in chemical gradients

With a quantitative understanding of how chemosensory diversity is affected by adaptation to different background stimuli, we wondered whether this property of the *K*_1/2_ distribution could have tangible effects on navigation. Conceptually, the impact of diversity tuning on navigation can be understood by considering a population climbing an exponential attractant gradient (**Figure 5A**). In such a gradient, the fold-change in stimulus concentration during a typical movement within the gradient is constant^40^. For a pure log-sensing system, this would mean the perceived change in signal is constant in the gradient, so if sensitivities to the gradient are diverse, they should remain diverse at all concentrations. However, for a system which displays diversity tuning, this is not always the case. At low ligand concentrations, cells will have diverse abilities to detect the gradient. Meanwhile, at high ligand concentrations, cells with different sensory capabilities will perceive similar signals. Our experiments suggest that the presence of other ligands in the environment should not affect this overall strategy, but can only, in the case of competitive ligands, change the concentration where this transition from diverse to near-uniform sensitivity occurs. We hypothesized that diversity tuning allows populations to benefit from high-performance cells at low concentrations, while attenuating the performance penalty of low-sensitivity cells as signal intensity increases.

**Figure 5.**
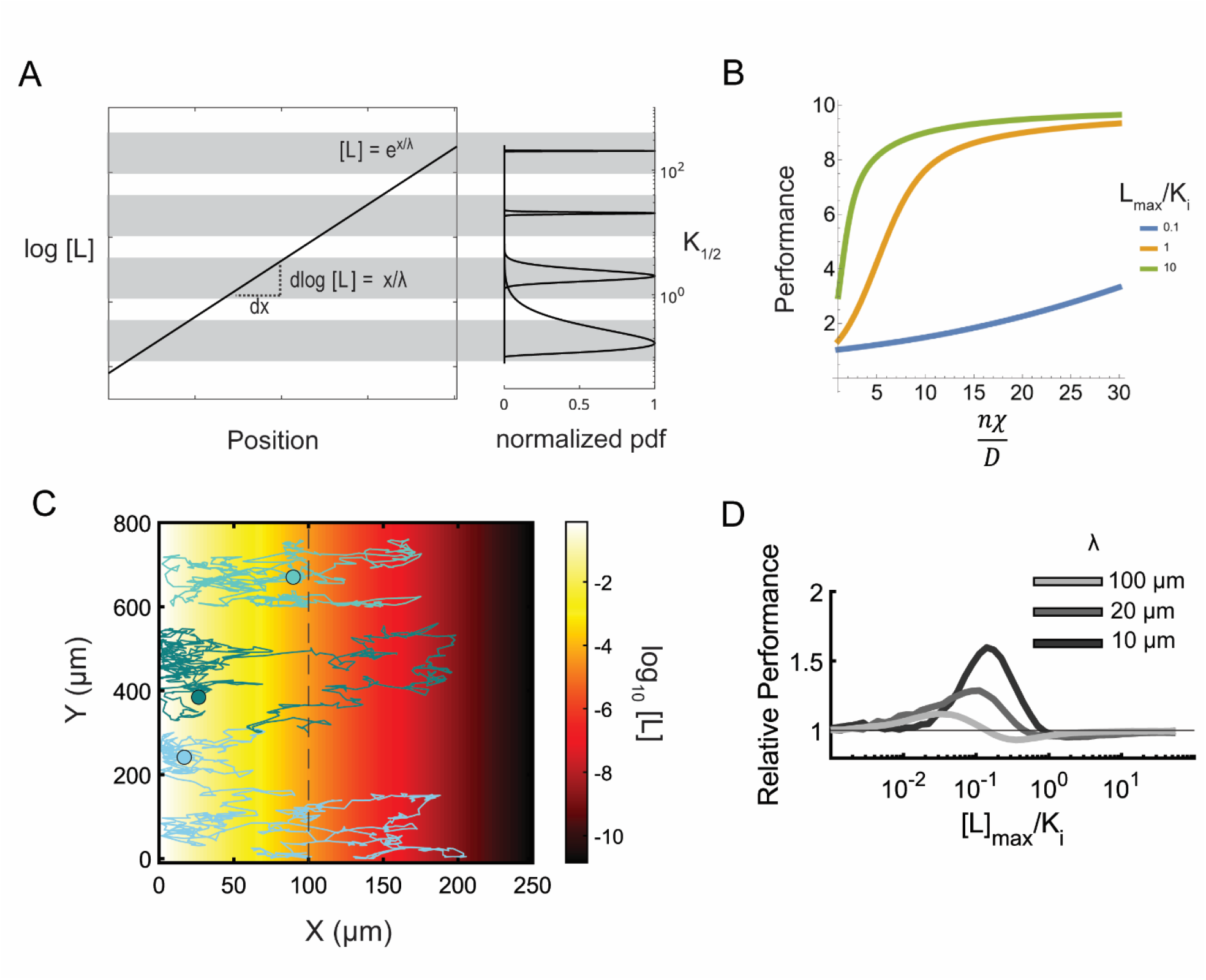
Diversity tuning during navigation in exponential attractant gradients. **A)** Schematic for the role of diversity tuning in navigation. (left) in an exponential gradient with length scale λ, the fold-change in stimulus concentration is constant in space. (right) Due to diversity tuning, how far in the gradient cells need to move to reach their *K*_1/2_ concentration varies at low concentrations, but is similar at high concentrations. As such, at different positions in the gradient, the impact of sensory diversity on navigation may vary. **B)** Analytical solution for the chemotactic performance – defined as the average ligand concentration experienced by a population with cooperativity *n*, chemotactic coefficient *χ*, and diffusion coefficient *D* – in a bounded exponential gradient with different maximal concentrations *L*_*max*_ at the bound. **C)** Agent-based simulations of cells navigating an exponential gradient. In the simulations, only receptor cooperativity is allowed to vary. All other phenotypic parameters are shared between individuals. The environment contains an exponential gradient of the ligand (log-concentration indicated by colorbar), with a reflective barrier at *x* = 0. Cells are initiated 100 µM from this barrier (dotted line) and allowed to navigate for 1500 seconds. Three example cell trajectories (colored lines) are shown with their final positions indicated by the filled circles. Trajectories are offset in the Y direction for clarity. **D)** Chemotactic performance of simulated agents was calculated using the final positions of 10,000 agents after 1500 seconds of simulation time. Relative performance is the performance of a population with diverse receptor cooperativity divided by the performance of a population where each cell has the median cooperativity. All other behavioral parameters were uniform in the population. The relative performance has a bilobed shape in gradients with a range of length scales. In shallow gradients (low concentration at the origin [*L*]_*max*_ relative to the dissociation constant *K*_*i*_), diverse cooperativities are advantageous as they allow rare high-performance cells to track the gradient. As signal strength increases, diversity incurs a mild performance deficit. In very strong gradients, the effect of diversity on performance is minimal.

To test this idea, we solved a mathematical model of chemotactic migration (STAR Methods) that takes into account the cooperativity of the receptor cluster^41,42^. This model is a simple Keller-Segel drift-diffusion equation, where the drift velocity is proportional to the perceived chemotactic signal. This perceived signal is calculated using the MWC model^41,43^, which, as we have seen above, exhibits sensory diversity tuning. Solutions were analytically tractable with exponential attractant gradients for a bounded arena with reflecting boundary conditions.

We defined the chemotactic performance of a phenotype as the average ligand concentration over the population density at steady state, relative to the average concentration over a density of cells incapable of sensing the gradient. Solving for this performance as a function of receptor cooperativity *n* revealed three performance regimes characterized by the concentration scale of the gradients (**Figure 5B**). At low concentrations, the chemotactic performance is a convex function of the cooperativity *n* over the entire range of *n* values. As the concentration increases, the performance curve enters a sigmoidal regime, and eventually a convex regime at high concentrations. These same concentration-dependent performance regimes were also found numerically for other gradient shapes including a Gaussian ligand profile (**Figure S5C**). These three regimes of the performance curve suggest that, due to Jensen’s inequality^44^, over any fixed interval of receptor cooperativities, the average performance of a population with diverse cooperativities can exceed the performance of the average phenotype in low concentration gradients, but may suffer a disadvantage when signals are strong.

While the qualitative effect of diversity on performance is predicted by the shapes of these performance curves, the actual magnitude of the advantage/disadvantage depends on the distribution of cooperativities, as well as the environmental stimulus field. To explicitly quantify such effects of diversity tuning on navigation performance, we utilized an agent-based simulation of chemotaxis to explore this effect in different environments with a distribution of sensory capabilities constrained by our measured *K*_1/2_ distributions. In the agent-based simulation, cells sense their environment using an MWC-type receptor cluster with cooperativity *n*. Receptor adaptation and flagellar motor behavior are simulated as previously described^45,46^ (STAR methods). We considered two populations: a diverse and a uniform population. In the diverse population, each cell’s cooperativity was sampled from a distribution approximated from the zero-background *K*_1/2_ distribution for MeAsp (**Figure S5A**). For simplicity, we assumed that all other phenotypic parameters did not vary. In the uniform population, each cell had the median cooperativity from this same distribution. In this way, we could compare the performance of a diverse population to the performance of the median phenotype.

For each population, we simulated 10,000 cells for 1500 seconds in three dimensions with an exponential gradient that decreased in one dimension, and a reflecting plane at *x* = 0 (**Figure 5C**). The concentration of the gradient reaches a maximum value [*L*]_*max*_ at this barrier. As in the analytical model, we defined the performance of a population as the average over the set of ligand concentrations corresponding to the set of final positions of cells in the population. We computed the performance of the diverse and uniform populations in attractant gradients with different maximal concentration in the environment, and various length scales. With all length scales tested, we found that when [*L*]_*max*_ was low, the performance of the diverse population exceeded that of the uniform population. As [*L*]_*max*_ increased, there was a regime in which the diverse population was at a slight disadvantage. At very high [*L*]_*max*_, the performances of both populations were similar (**Figure 5D**). In simulations within arenas with two separate Gaussian attractant peaks, we observed qualitatively similar performance curves when the attractants bound uncompetitively, and competitively (**Figure S5D-H**).

Both the analytical model and agent-based simulations demonstrate that diversity tuning can play a role in chemotactic navigation. Variation in phenotypic parameters will generate cells with variable chemotactic ability. However, diversity tuning allows the variation in chemotactic ability to change with the environment without changes in the underlying phenotype distribution. When signals are weak, the convex phenotype-performance map leads to a regime where performances are variable and the population can benefit from rare high-sensitivity cells. When signals are very strong, the phenotype-performance map flattens, leading to similar performance of all phenotypes.

### Diversity tuning beyond chemotaxis

Given the navigational performance benefits suggested by our mathematical analysis and simulations above, we next asked if analogous diversity tuning arises in more general adaptive signaling systems beyond bacterial chemotaxis and the allosteric MWC model. In the STAR Methods, we present a mathematical analysis that identifies a general class of sensory systems, which includes the MWC model of chemotaxis, in which background-dependent sensory diversity tuning arises. In particular, we demonstrate that a sufficient condition for sensory diversity tuning is that the sensory system in question demonstrates upon adaption to a background input level *L*_0_ a dose-response relation that can be written as a function

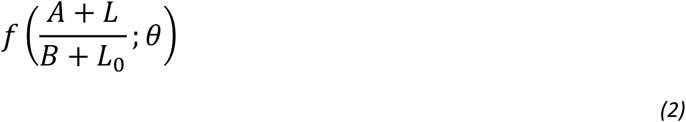

where *f* is a monotonic invertible function with phenotypic parameters *θ* which vary from cell to cell, *A* and *B* are positive constants analogous to dissociation constants, and *L* is the foreground input level. This generic dose-response relation, which could arise from a variety of underlying mechanisms, demonstrates two sensory regimes. When *L*_0_ is low relative to *B*, the system senses linear changes in ligand concentration. However, when *L*_0_ greatly exceeds *B* the system responds to the fold-change in ligand concentration relative to the background. For systems with such dose-response curves, we define a sensitivity *K*_*R*_ analogous to the *K*_1/2_ but for arbitrary response level *R*. The coefficient of variation in *K*_*R*_ can be shown in general to decrease as the background ligand concentration increases. Furthermore, if *A* = *B*, the system is now precisely adapting over all background stimuli, and the CV of *K*_*R*_ is exactly equation (1) in the absence of a competitive background species (STAR Methods). As an illustrative example, we show how a dose-response curve in the form of equation (2) can arise in simple adaptive feed-forward and feed-back networks in the STAR Methods.

While equation (2) is a sufficient condition for tunable diversity, the dynamic range of the sensory diversity (i.e. the maximal achievable contrast in the *K*_*R*_ CV) does depend on the specific form of *f*. Specifically, fractional change in sensory diversity as the population adapts to large backgrounds depends on the average zero-background sensitivity relative to the dissociation constant *A* (STAR Methods). In the case of chemotaxis, what this implies is that if cells are on average more sensitive than individual receptors (*K*_0_ < *K*_*i*_), then sensory diversity will change significantly as the population adapts to large background stimuli. However, if cells are not more sensitive than individual receptors (*K*_0_ ≈ *K*_*i*_) the change in diversity as cells adapt will be small. In the case of *E. coli* chemotaxis, high sensitivity is accomplished by cooperativity in the receptor cluster. This suggests that both the transition from linear to fold-change detection, and high cooperativity may be important for sensory diversity tuning.

## Discussion

With decades of extensive characterization, *E. coli* chemotaxis provides an excellent testing ground for general principles of signal processing by living systems, and the role of cell-to-cell variation in population-level sensory strategies. In this study, we used single-cell FRET to measure the sensitivity distributions for several ligands after adaptation to different background stimuli. Consistent with previous studies, increasing the concentration of a particular ligand led to a decrease in the amount of *K*_1/2_ diversity for that ligand.^27^ Here, we found that if two ligands bind uncompetitively – on the same receptor or different receptors – sensory diversity for each ligand can be tuned in an orthogonal manner. That is, it is possible to be in the low-diversity regime for one ligand, and the high diversity regime for another ligand in the same environment. However, for ligands that compete for binding sites, increasing the background concentration of one ligand decreases *K*_1/2_ diversity for that ligand, while increasing the *K*_1/2_ diversity of its competitors.

Previously, we showed that the two diversity regimes are explained by the two regimes of allosteric receptor clusters. At low methylation levels, the *K*_1/2_ is strongly dependent on the receptor cooperativity. However, when receptors are highly methylated, the *K*_1/2_ depends weakly on the receptor cooperativity^27,35^. Our experiments and theory suggest that what matters for sensory diversity tuning is not simply the methylation level of the receptors, but the amount of receptor methylation that directly offsets the free energy contribution from each ligand species.

First, consider two ligands that bind uncompetitively such as serine and MeAsp. Increasing the background MeAsp concentration leads to receptor methylation to offset the effect of MeAsp on the kinase activity. The magnitude of this offset, as in the single species case, determines whether the population is in the high or low diversity regime.^27,35^ Adding serine to the background will cause additional receptor methylations that offset the serine concentration. If we change the MeAsp concentration however, responses will not be affected by these new methylations, since their effects are canceled out by the constant serine background. As such, the distribution of MeAsp sensitivities remains independent of the serine concentration. Only methylation due to the background MeAsp concentration affects the MeAsp *K*_1/2_ distribution.

The case of competing ligands is slightly more complicated. Consider MeAsp and L-Asp for example. Again, adapting to MeAsp leads to receptor methylations which offset the background MeAsp concentration. However, addition of a competitive background ligand, while leading to additional methylations, now also changes the effective affinity of MeAsp for the receptor due to reduced binding site availability. Due to this change in the effective affinity of the receptor for MeAsp, even though the total receptor methylation has increased, the total contribution of MeAsp to the free energy has decreased, and as such a smaller proportion of the total receptor methylation is offsetting its concentration. This decrease in methylation required to offset its concentration leads to an increase in the MeAsp *K*_1/2_ diversity.

Another, more general way, to understand sensory diversity tuning is in terms of the transition from a linear sensing regime, where cells sense the total change in ligand concentration, to a logarithmic sensing regime, where cells sense the fold-change in ligand concentration relative to the adapted background concentration. In bacterial chemotaxis, the implementation of this transition (see equation (2)) by receptor clusters as described above is sufficient to capture the *K*_1/2_ diversity as background conditions change. From this perspective, the reason for which sensory diversity can be tuned orthogonally for ligands that do not compete for binding sites is that the transition from linear to logarithmic sensing is also orthogonal for such ligands. However, for competitive ligands, previous theory showed that only one ligand species can be in the logarithmic sensing regime at a time.^47^ While there are likely milder conditions for diversity tuning than equation (2), dose-response relationships of this kind can emerge in some simple feed-forward and feed-back signaling pathways (STAR Methods),^40,48^ suggesting that diversity tuning may arise in adaptive signaling systems beyond chemotaxis.

Functionally, how might diversity in sensitivity (*K*_1/2_) and the ability to tune this diversity in different backgrounds benefit bacterial populations? Behavioral studies previously revealed diversity in the chemotactic gain, and suggested that maintaining poor navigators in a population can protect the population from catastrophic events.^19^ Additionally, within each cell, temporal fluctuations in gain have been theorized to improve resource exploitation in complex environments by preventing cells from getting trapped in local attractant maxima.^17^ One implementation of pathway gain diversity which is consistent with our model, may be variation in receptor cooperativity, which has been suggested by previous work.^19,27^

Our findings add context to these previous results. Variation in receptor cooperativity can generate rare high-sensitivity cells that can track weak signals. But as the signal prevalence increases, diversity tuning enables even rare low-sensitivity cells to track the same gradient. This perspective is supported by our mathematical modeling of chemotactic behavior where we challenged cells with diverse sensory capabilities to track an exponential gradient, or locate the center of two different gaussian attractant signals. When the attractant source is weak, a diverse population has higher performance than the median phenotype since rare high-gain cells can track the gradient. However, at high source strength, due to diversity tuning, the low-gain cells do not significantly hamper performance, and behavior is similar to that of the median phenotype.

Ultimately, our experiments and theory suggest that *E. coli* implement sensory diversity tuning in a manner that allows the population to simultaneously deploy a bet-hedging strategy for responding to some ligands while deploying a tracking strategy to climb gradients of others. This feature of the chemosensory receptor cluster could aid populations in making behavioral decisions. If weak signals appear, a small segment of the population will track them without committing the entire population to a potentially transient signal. Due to precise adaptation and the independence of the sensory diversity for different ligands, this behavioral strategy can be applied independently to different signals by the same population, at the same time.

## Acknowledgements

We thank JS Parkinson and V. Sourjik for providing strains, and we thank F. Avgidis, D. Clark, N. Dimitrova, and H. Mattingly for useful discussions.

## Author contributions

J.M., K.K. T.E., and T.S.S. designed the project. K.K. designed the experimental setup. J.M. and R.K. performed the experiments. J.M. performed the data analysis and mathematical modeling. J.M., K.K., R.K., T.E. and T.S.S. wrote the manuscript.

## Data and materials availability

All data needed to evaluate the conclusions in the paper are present in the paper and/or the Supplementary Materials. Additional data may be requested from the authors.

## STAR Methods

### RESOURCE AVAILABILITY

#### Lead contact

Further information and requests for resources should be directed to and will be fulfilled by the Lead Contact, Thierry Emonet.

#### Materials availability

This study did not generate new unique reagents or strains.

#### Data and code availability

All data needed to evaluate the conclusions in the paper are present in the main text and/or the supplementary materials. The datasets generated in this study have been deposited in the dryad database (doi: 10.5061/dryad.nvx0k6dzz). Codes used in this study are available on GitHub (https://github.com/emonetlab/SensoryDiversityTuningAnalysis). Any additional information required to reanalyze the data reported in this paper is available from the lead contact upon request.

### EXPERIMENTAL MODEL DETAILS

#### Bacterial growth conditions

All FRET experiments were performed with a derivative of *E. coli* K-12 RP437 harboring ΔCheYZ and ΔFliC mutations, and transformed with two plasmids: pSJAB106 from which CheZ-YFP and CheY-mRFP1 are expressed in tandem on an isopropyl-β-d-thiogalactopyranoside (IPTG) inducible promoter, and pZR1 from which ‘sticky’ FliC* is expressed on a sodium-salicylate (NaSal) inducible promoter.^21^ Cells were grown overnight in TB broth (1% bacto-tryptone, 0.5% NaCl), then diluted 1:100 into 10mL of fresh TB supplemented with 50*µM* IPTG to induce the FRET pair, and 3*µM* NaSal to induce sticky FliC* for cell adhesion to glass coverslips^27^, with 100ug/mL ampicillin and 34ug/mL chloramphenicol for plasmid retention. Cultures were grown at 33.5°*C* to an OD_600_ of 0.44-0.47 then washed once with 30mL motility buffer (10mM KPO4, 0.1mM EDTA, 1uM methionine, 10mM lactic acid) and suspended in 2mL of motility media. Suspensions were left at room temperature for 2hrs before imaging.

### METHOD DETAILS

#### Microfluidics fabrication and operation

Microfluidic devices were constructed from PDMS with standard soft lithography methods.^49^ The master mold for the device was a silicon wafer with features created using ultraviolet photoresist lithography (Figure S1). To cast the device, the mold was coated with a 5 mm-thick layer of degassed 10:1 PDMS-to-curing agent mixture (Sylgard 184, Dow Chemical). Molds were baked at 80°C for 45 min. Devices were cut out of the mold and holes were punched into the inlets with a manually sharpened 20-guage blunt-tip needle. Then, PDMS devices were bonded to a 24mm X 60mm coverslip (#1.5). PDMS was cleaned with transparent adhesive tape (Magic tape, Scotch) then rinsed with acetone, isopropanol, methanol, and water. Coverslips were also rinsed with acetone, isopropanol, methanol, and water. Glass and PDMS were treated with plasma generated by a corona treater. Then, PDMS was laminated to the coverslip, and baked overnight at 80°C.

The microfluidic device has 7 inlets, an outlet at the end of the imaging chamber, and two side outlets of which only the right one is opened during cell loading (Figure S1). During operation, the inlets were connected to 7 reservoirs containing motility media supplemented with different attractant concentrations. Cells washed in motility medium were loaded into the device through the outlet channel, where they initially exit through one of the side outlets, which is plugged before imaging to induce flow towards the end outlet. Pressure in the inlet reservoirs was controlled by computer-controlled solenoid valves (MH1, Festo) that quickly switch between atmospheric pressure, and a source of 1kPa pressurized air as needed.

#### In-vivo single-cell FRET microscopy

Single-cell FRET and cell culture were performed as previously described.^27^ Cells were grown and washed as described above. FRET imaging was performed with an inverted microscope (Eclipse Ti-E, Nikon) equipped with an oil immersion objective (CFI Apo TIRF 60X Oil, Nikon). Yellow fluorescent proteins were illuminated with a light-emitting diode system (SOLA SE, Lumencore) through one excitation filter (59026x, Chroma), then another (FF01-500/24-25, Semrock) and a dichroic mirror (FF520-Di02; Semrock). Emission was fed into an emission image splitter (OptoSplit II, Cairn) where it was split into donor and acceptor channels with a dichroic mirror (FF580-FDi01-25×36, Semrock) and collected through emission filters (FF01-542/27 and FF02-641/75; Semrock) with a scientific CMOS camera (ORCA-Flash 4.0 V2, Hammatsu). Red fluorescent protein mRFP1 was imaged in the same way as YFP, except with a different second excitation filter (FF01-575/05-25) and dichroic mirror (FF593-Di03-25×36, Semrock). For both fluorophores, images were taken with 50 ms exposure time.

#### FRET data analysis

Single-cell fluorescent signals were extracted from fluorescence time-series as described previously.^21,27,50^ Images were segmented and single-cell fluorescent signals determined with in-house software. Photobleaching was corrected by fitting donor and acceptor time-series with a bi-exponential function and subtracting out the decay to yield donor *D*(*t*) and acceptor *A*(*t*) time-series.

To calculate FRET from fluorescence time-series, we used the E-FRET method.^50^ The E-FRET index is given by

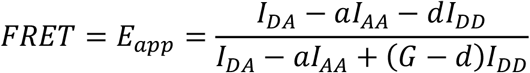

where *I*_*DA*_ is FRET-acceptor emission intensity from donor excitation, *I*_*AA*_ is acceptor emission with acceptor excitation, *I*_*DD*_ is donor emission from donor excitation, with *a, G, d* being optical constants that depend on the FRET pair and optical setup which were determined by an independent experiment with strains that express only CheY-mRFP or only CheZ-mYFP.^22^

To convert FRET time-series to kinase activity, for each cell, FRET values were normalized by the minimum and maximum values measured during a saturating stimulus at the beginning and end of the experiment. For each experiment, the saturating stimulus was a mixture of the background stimulus, and a saturating amount of the foreground stimulus.

#### Determining K_1/2_ distribution from single-cell responses

After a saturating stimulus for response calibration, cells were adapted to the background for 3 minutes. Then, cells were presented with 5 different attractant concentrations in series with 5 second presentation time, and 30 seconds at the background level in between stimuli. This cycle through the stimulus levels was presented 7 times to allow within-cell averaging of the response amplitudes.

We extracted the *K*_1/2_ cumulative distribution function from the within-cell average responses using previous methods.^27^ Given a family of dose-response curves, for a given ligand concentration [*L*]_*i*_, the fraction of responses that are greater than half-maximal equals the fraction of cells with *K*_1/2_ < [*L*]_*i*_ (Figure S1). Thus, by determining the fraction of half-maximal responses at a series of ligand concentrations, we can measure the cumulative distribution function (CDF) of the *K*_1/2_. For each measured point on the CDF, confidence intervals were determined by bootstrapping, both over cells and over the seven responses per re-sampled cell. Confidence intervals (95% CI) on the fitted lognormal parameters were estimated from the posterior distribution with lognormal priors and gaussian error using markov-chain monte carlo methods (Figure S2). For each CDF, cells from at least two independent biological replicates were pooled to increase the number of cells included in each analysis, since the number of cells measured in each experiment varied from day to day.

#### Model of the chemotaxis pathway

Throughout the main text and STAR Methods, we use the standard model of bacterial chemotaxis^32-35,51^ where the cell relays sensory information to the flagellar motor through a signaling cascade triggered by binding of attractant to highly cooperative receptor arrays. Receptor clusters whose activity can be described by a two-state model where activity *a* ∈ [0,1] is determined by the free energy difference *F* (in units of *k*_*B*_*T*) between the active and inactive states

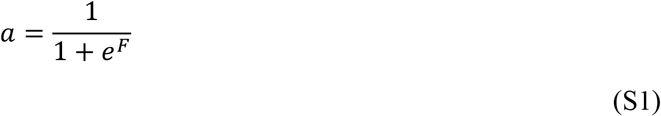

The free energy can be decomposed into two terms 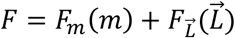. The first depends linearly on the degree of receptor methylation *m*, such that *F*_*m*_(*m*) = −*n*_*T*_α(*m* − *m*_0_) where *n*_*T*_ is the total receptor coupling, and α and *m*_0_ are constants.^52^ The second term is the dependence of the free energy on ligand binding, where 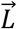 is a vector containing the concentrations of all ligand species in the environment. The form of *F*_*L*_ (*L*) will depend on whether the ligands bind competitively or uncompetitively to the receptor cluster. For a receptor cluster with two receptor species - Tar and Tsr - that are allosterically coupled, and two ligands that bind either Tar or Tsr but not both,

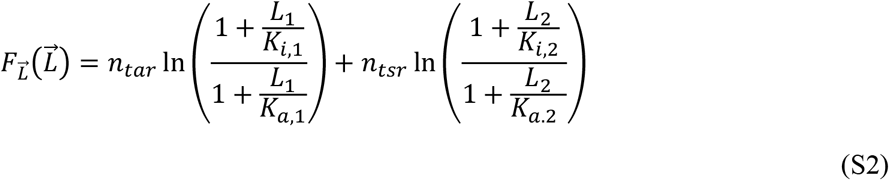

where *n*_*tar*_ and *n*_*tsr*_ are the number of cooperative Tar and Tsr units in the cluster with *n*_*tar*_ + *n*_*tsr*_ = *n*_*T*_, *L*_1_ and *L*_2_ are concentrations of ligands binding Tar and Tsr respectively with dissociation constants *K*_*i*_ and *K*_*a*_ to the inactive and active receptor states. The case where two ligands bind independently to the Tar receptor is identical to equation (S2) with *n*_*tsr*_ = *n*_*tar*_. In cases where *L*_1_ and *L*_2_ both bind Tar in the same binding pocket,

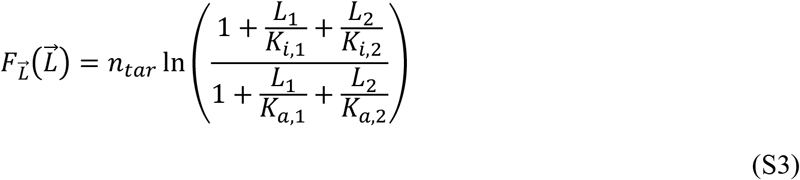

As the external ligand concentration changes, receptors are methylated or demethylated to adapt to maintain an activity level *a*_0_ independent of the environment. To describe methylation kinetics, we used a similar approach as previous studies, where

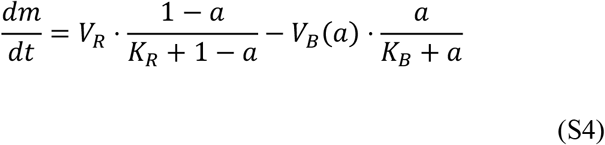

where *V* and 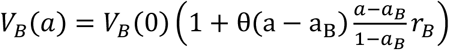 (*θ* is the Heaviside function and *a*_B_ = 0.74, *r* = 4.0) are the methylation/demethylation rates for CheR and CheB with dissociation constants, *K*_*R*_ and *K*_*B*_.^18,33,52^

We assume signal transduction to be fast compared to methylation, so that the active response regulator CheY-P concentration can be written, *Y*(*a*) = α_*y*_*a* with α_*y*_ constant. CheY-P binds the flagellar motor, changing the switching rate between clockwise (CW) and counter-clockwise (CCW) rotation. This switching is assumed to be a Poisson process with switching rates

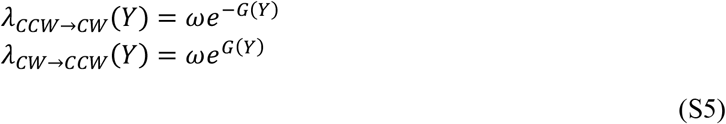

Where 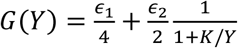 with *ϵ*_1_, *ϵ*_2_, *K* constant.^18,45,53^ We assume that only a single motor switching from *CCW* → *CW* is sufficient to tumble, so for cells with only 1 flagellum, the rates above describe the run and tumble dynamics as a function of CheY-P.

#### Agent-based simulations of chemotactic behavior

Agent-based simulations were performed using Euler integration of the above model, as described previously.^18,45,53,54^ At each time-step, the cell either moves or stays in place depending on its motility state (run or tumble), which both have their own rotational diffusion coefficients *D*_*R*_ and *D*_*T*_. After updating the position and local ligand concentration, the adaptation equation is integrated and the free energy of the receptors, and thus the CheY-P concentration and motor switching rates are updated. A random number is drawn to determine whether the flagellum state switches, with rules and parameters as in.^53^

Cells were challenged to navigate two different environments. The first is a simple exponential gradient extending in one dimension with a reflecting boundary at *x* = 0, maximum concentration *L*_*max*_, and length scale λ, given by *L*(*x*) = *L*_*max*_*e*^−*x*/λ^. The second was an environment with two static gaussian gradients, given by 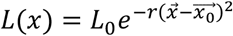, where *L*_0_ is the ligand concentration at the starting position, 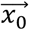 is the center of the gaussian, and *r* is the gradient ramp rate. The gradients were placed equidistant from the origin where cells were initialized. Additionally, for each cell, values of *n*_*tar*_ and *n*_*tsr*_ were drawn by sampling from the experimental *K*_1/2_ distribution for meAsp in 0-background, and using the conversion 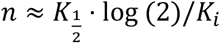.^35^ For simplicity, the same distribution was assumed for both *n*_*tar*_ and *n*_*tsr*_. Additionally, the distribution of *n*_*tar*_ and *n*_*tsr*_ was assumed to be independent since responses to the population average *K*_1/2_ were uncorrelated (Figure S5B). To isolate the effect of signal integration and diversity in receptor coupling, we assumed that methylation does not saturate, which would lead to imperfect adaptation as the population climbs the gradient.^55,56^

#### Quantitative model of the K_1/2_ CV as a function of background ligands

Here we derive the coefficient of variation (CV) of the distribution of *K*_1/2_ for the classic MWC model of bacterial chemoreceptor clusters. For a ligand L, assuming *L* ≪ *K*_*a*_, the activity of the cluster is approximately

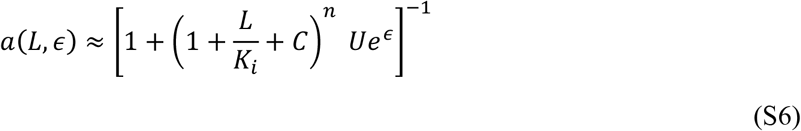

where *ϵ* is the free energy of the cluster in the absence of all ligands, *C* = *∑*_*l*_ *L*_*l*_/*K*_*l*_ is a constant that accounts for the presence of all background attractants that compete with L, and *U* is a constant that accounts for all uncompetitive ligands. *C* and *U* follow from equations (S2) and (S3) above. Note that *U* can take different forms depending on how the uncompetitive ligand binds. For example, if the second ligand *L*′ bound the same receptor as *L* with dissociation constant 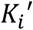, then 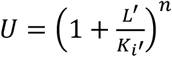. However, if it bound a different receptor which contributed *n*^′^ cooperative units to the receptor cluster, then 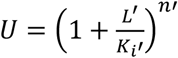.

We assume that the receptor cluster adapts perfectly. This means that at steady state, any combination of ligands will result in the same activity, given by

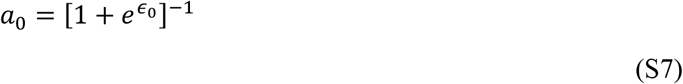

where *ϵ*_0_ is the adapted zero-background free energy difference. Now, we can define the *K*_1/2_ as the ligand concentration *L* where *a*(*L, ϵ*) = *a*_0_/2. We also write *L* = *L*_0_ + Δ*L* where *L*_0_ is the background concentration of *L*. To find the *K*_1/2_, we solve

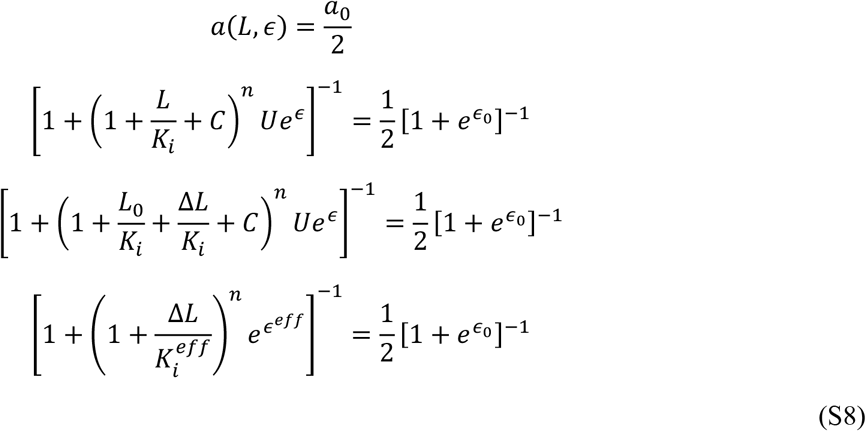

Where 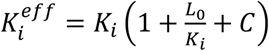 and *ϵ*^*eff*^ = *ϵ* + In(*U*) + *n* ⋅ In(1 + *C* + *L* /*K*) = *ϵ* due to perfect adaptation. Continuing,

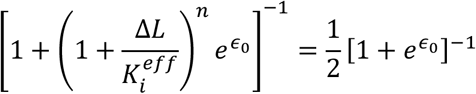

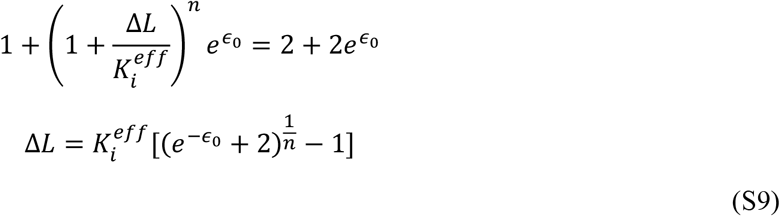

Thus, the *K*_1/2_ as defined above, is given by

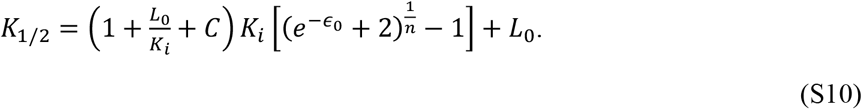

In the zero-background condition, the sensitivity becomes 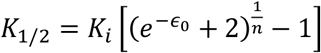

The *K*_1/2_ clearly depends on both the free energy of the cluster in zero background condition *ϵ*_0_, and the degree of receptor coupling, *n*. It also depends on the concentration *L*_0_ of ligand in the background, as well as on the presence of competitive ligand *C* = *∑*_*l*_ *L*_*l*_/*K*_*l*_ in the background. However, the *K*_1/2_ is unaffected by the presence of uncompetitive ligands, whether the bind the same or different receptors. We assume that the receptor-ligand affinities *K*_*i*_ and *K*_*l*_ and concentrations *L*_0_ and *L*_*l*_ of the ligand and competitors in the background are the same for each cell. Thus, we can write the average *K*_1/2_ as

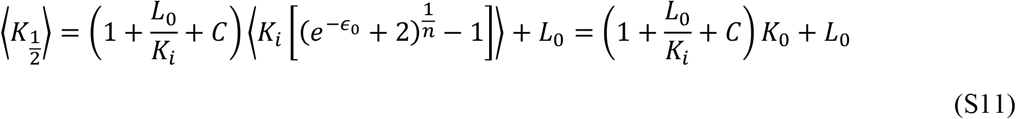

and the standard deviation as

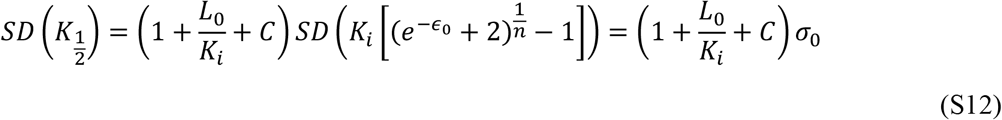

Where 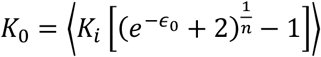 and 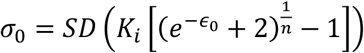 are equivalent to the mean and standard deviation of the *K*_1/2_ distribution for ligand L in the absence of any background stimulus. Then, the coefficient of variation (*CV* = *SD*(*K*_1/2_)/⟨*K*_1/2_⟩) in *K*_1/2_ for ligand L in the presence of a background concentrations *L*_0_ of itself and background concentrations *L*_*l*_ of competitors is given by

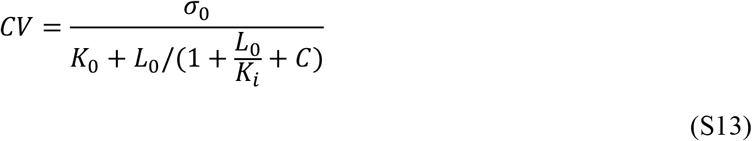

By inspection and for increasing *L*_0_, the maximal CV is *σ*_0_/*K*_0_ and the minimal CV is *σ*_0_/(*K*_0_ + *K*_*i*_). Given a particular competitive background *C*, the transition between the high and low diversity regimes as *L*_0_ increases occurs when

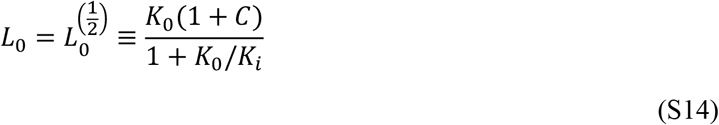

#### Diversity tuning emerges from the transition from linear to logarithmic sensing

The molecular details of the chemosensory network are not necessary to capture the essential diversity-tuning behavior of the chemotaxis pathway. A sufficient condition to have background dependent sensory diversity is that the dose-response relationship can be written as a function

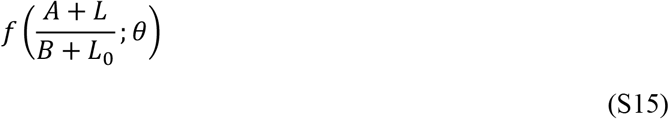

where *A* and *B* are positive constants, *L*_0_ is the background stimulus level, *L* is the foreground stimulus level, and *f* some monotonic invertible function with phenotypic parameters *θ*. Note that this function is always precisely adapting if *A* = *B*, but otherwise only precisely adapting when *L*_0_ ≫ *A, B*. Analogous to the *K*_1/2_, the stimulus level where activity is half of the steady state, we define a *K*_*R*_, which is the stimulus level where 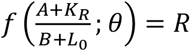. We define the inverse of the monotonic function *f* at constant *θ* by 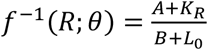. Solving for *K*_*R*_,

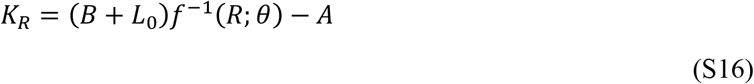

We define *K*_0_ = *B* ⋅ ⟨*f*^−1^(*R*; *θ*)⟩_*θ*_ − *A*, which is the mean of the distribution of *K*_*R*_ in zero-background (*L*_0_ = 0), and the subscript *θ* denotes an average over the distribution of phenotypic parameters *θ*. Similarly, we define *σ*_0_ = *B* ⋅ *SD*_*θ*_(*f*^−1^(*R*; *θ*)) which is the zero-background standard deviation of the distribution of *K*_*R*_, and *SD*_*θ*_ is the standard deviation of the distribution of *θ*. In an arbitrary background, the 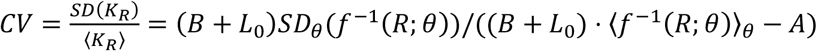. Using the definition of *K*_0_ and *σ*_0_ to replace ⟨*f*^−1^(*R*; *θ*)⟩_*θ*_ and *SD*_*θ*_(*f*^−1^(*R*; *θ*)), we obtain

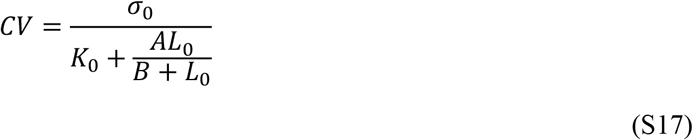

which is a decreasing function of the background ligand concentration *L*_0_. If *A* = *B* = *K*_*i*_, equation (S17) is identical to equation (1) from the main text in the absence of competitive background ligands.

The stated sufficient condition guarantees that the degree of cell-to-cell variability will vary with the background stimulus level. But is the change in variability always significant? It can be shown that the difference between the maximum and minimum levels of cell-to-cell variability depend only on the average sensitivity *K*_0_, relative to the dissociation constant *A*. As written above, the maximum *CV* occurs at *L*_0_ = 0, and the minimum *CV* occurs as *L*_0_ → ∞. These are,

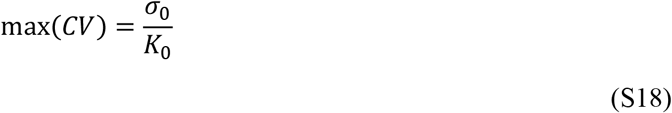

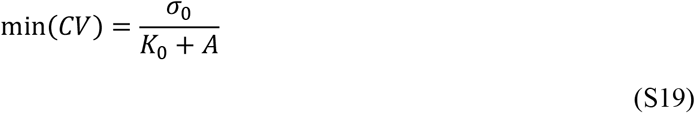

To determine the relative difference between the maximum and minimum *CV* we compute 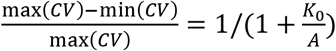 which reveals that the contrast between the high and low diversity states (i.e. the significance of diversity tuning) depends only on *K*_0_/*A*, the average zero-background *K*_*R*_ relative to the dissociation constant *A*. This leads to an intuitive understanding of the conditions for diversity tuning within this framework. If the cells achieve a sensitivity greater than the dissociation constant (*K*_0_ ≪ *A*), then adaptation can dramatically affect the *K*_*R*_ distribution. However, if cells are far less sensitive than ligand binding alone (*K*_0_ ≫ *A*), then adaptation has a negligible affect on diversity.

It is easy to check that this theory applies to diversity tuning in chemotaxis. We start by re-writing the MWC model in the form of Equation (S15). Assuming *L* ≪ *K*_*a*_, the activity of the receptor cluster can be written

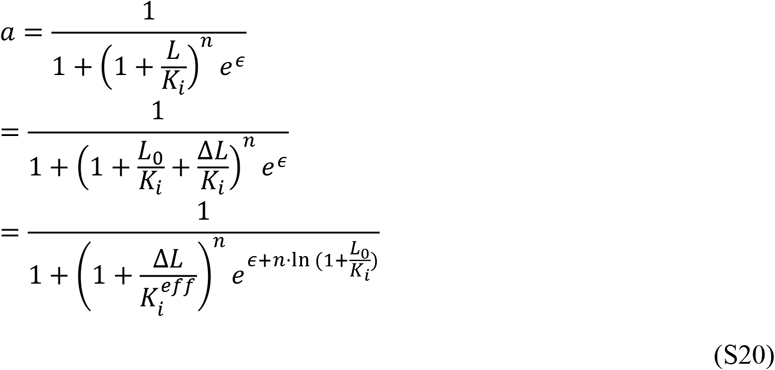

where, as before, we have decomposed *L* into its background concentration *L*_0_ and the change in foreground concentration Δ*L*, and defined 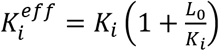. Assuming the activity adapts precisely to *L*_0_, *ϵ* will adjust until the sum 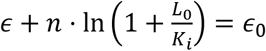 is the steady-state free energy difference. From here we get

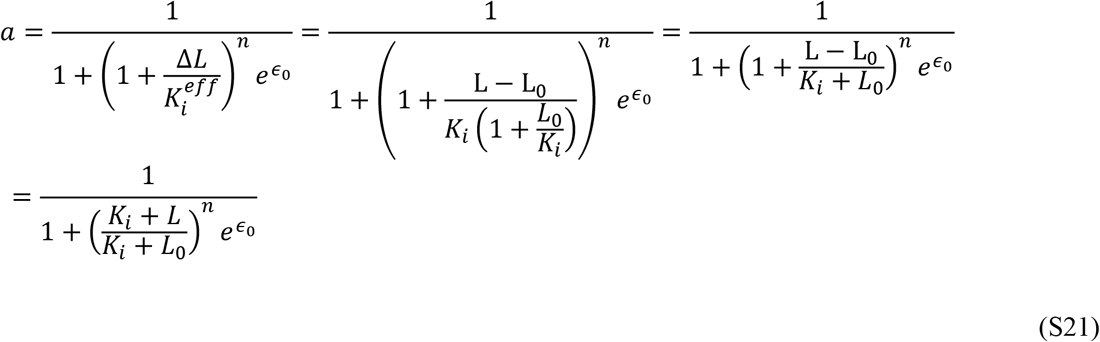

Therefore we have written the kinase activity *a* as a function of the form 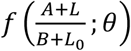 with *A* = *B* = *K*_*i*_ :

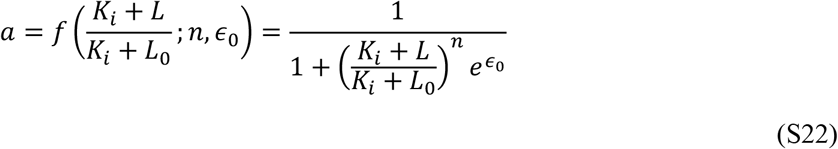

Given an activity level *a* = *r* the inverse of the function *f* at constant parameters *θ* = (*n, ϵ*_0_) is

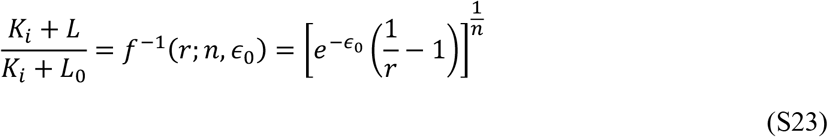

which can be used to calculate the ligand concentration *L* = *K*_*R*_ needed to reach the activity *a* = *r*:

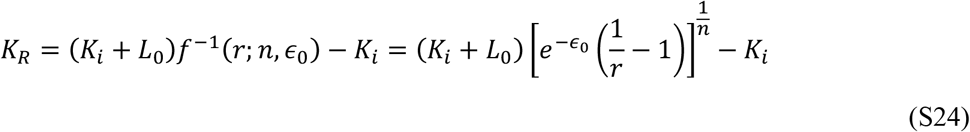

Here we are interested in the concentration *L* = *K*_1/2_ at which the activity is half its adapted value 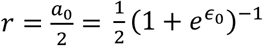. Inserting in the above expression we recover the expression (S10) for the *K*_1/2_ derived earlier.

#### Two simple examples of adaptive signaling pathways with diversity tuning

To illustrate that diversity tuning can arise when equation (S15) is satisfied in systems beyond chemotaxis, we analyzed two simple adaptive signaling pathways. With mild assumptions, diversity tuning can be shown to arise in two basic adaptive network motifs: the incoherent feed-forward loop, and negative integral feedback. First, we will consider a simple incoherent feed-forward loop with three species: the output species *Z*, the input *X* which activates *Z*, and the adaptation species *Y* which deactivates *Z* but is itself activated by *X*. We can write a zero-order approximation for the *Z* and *Y* dynamics^40,57-59^

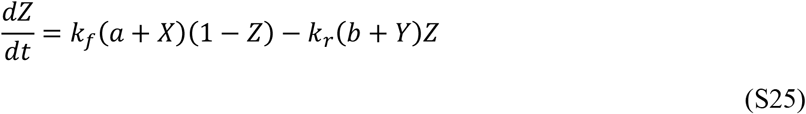

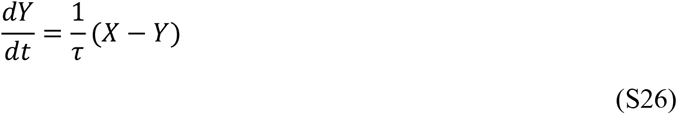

where *k*_*f*_ and *k*_*r*_ are rate constants, *a* and *b* are constants allowing constant basal production and destruction of *Z*, and τ is the timescale of *Y* equilibration. After adaptation to a constant background input *X*_0_, *Y* = *X*_0_.

Assuming adaptation is slow relative to the response of *Z* to change in *X*, the dose-response curve of *Z* activity is

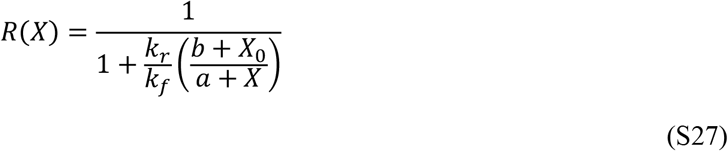

which is a function of the form of equation (S15), suggesting this system is capable of diversity tuning if *k*_*r*_ and/or *k*_*f*_ vary from cell to cell.

We can perform a similar analysis for negative feedback. For this case, we consider the same dynamics of *Z*, but different dynamics for *Y*. The negative feedback loop equations are thus

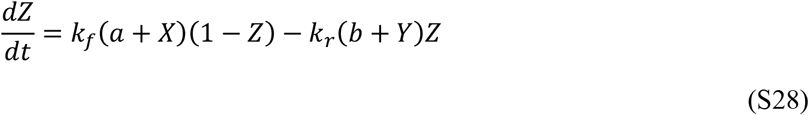

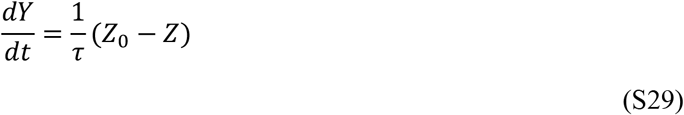

At steady state, assuming constant input *X*_0_, *Z* = *Z*_0_, so 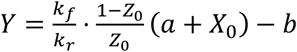, and the steady-state dose-response curve of Z is

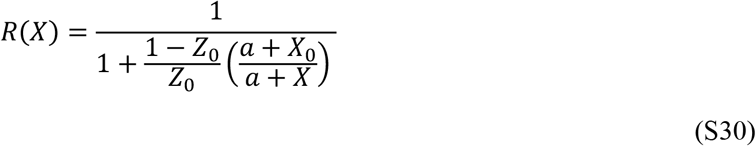

which is also a function in the form of equation (S15), and should also tune diversity due to variation in *Z*_0_ as the background input *X*_0_ changes.

#### Calculating chemotactic performance with diverse receptor cooperativity

To explore analytically the role of diverse receptor cooperativity in navigation, we used an extension of the classic Keller-Segel model.^41,42^ In this model, the bacterial cell density *ρ*(*x, t*) is governed by the partial differential equation

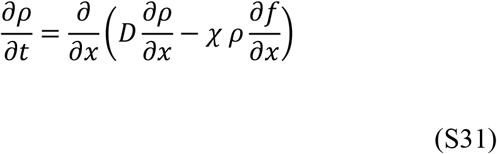

where *D* is the diffusion coefficient, *χ* is the chemotactic coefficient, and *f* is the signal perceived by the cell. This perceived signal is the change in free energy of the chemoreceptor complex given a change in ligand concentration, which as in the MWC model can be expressed as

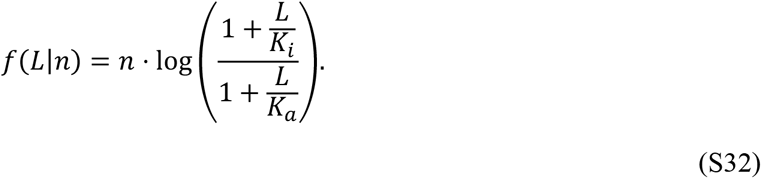

Assume a fixed ligand profile over the domain 0 ≤ *x* ≤ *a* and that *L*(*x*) ≪ *K*_*a*_. Then the cell density reaches a phenotype dependent steady-state

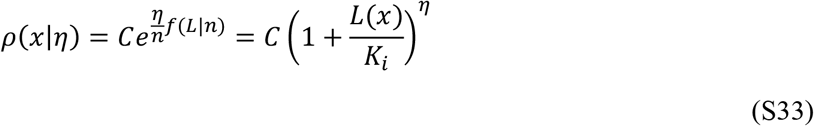

where *C* is the integration constant and *η* = *nχ*/*D*. It is convenient to define a conditional probability density of the cell position given phenotype *η* :

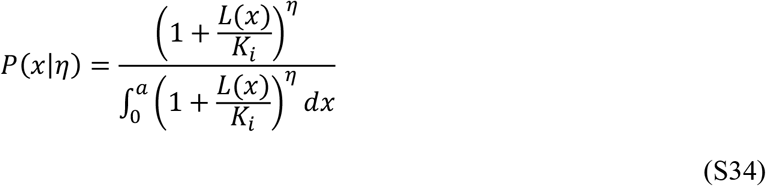

The cell density for phenotype *η* is then simply *ρ*(*x*|*η*) = *N*_*cells*_*P*(*x*|*η*). We also see that when cells do not respond to the gradient or when they do respond but there is no gradient, i.e. *L*(*x*) = *L*_0_ is constant, then the conditional probability density of the cell position becomes a constant *P*_0_ :

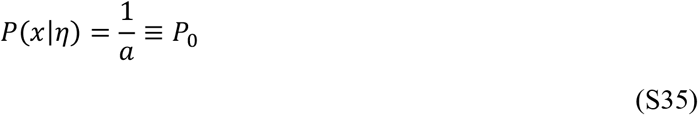

In this case the cell density is uniform and becomes *ρ* = *N*_*t*°*t*_*P*_0_ = *N*_*t*°*t*_/*a*, as expected.

We now define the performance of a particular phenotype as the expected ligand concentration experienced by the cells with phenotype *η* normalized by expected ligand concentration experienced by the cells when they are not responsive to the ligand:

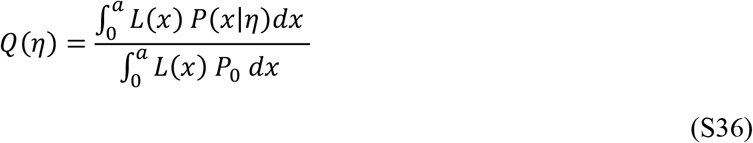

For an exponential gradient *L*(*x*) = *L*_*max*_*e*^−*x*/λ^ we calculate *P*(*x*|*η*) by changing variables from *x* to

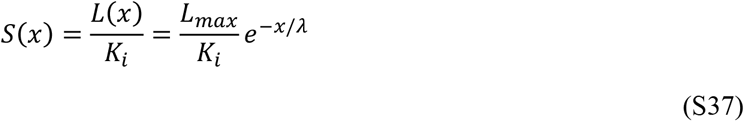

Introducing (S37) in equation (S34) using the fact that 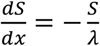 we get:

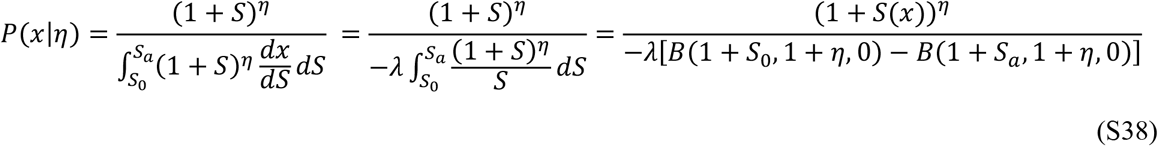

where *S*_0_ = *S*(0), *S*_*a*_ = *S*(*a*) and *B* is the incomplete Beta function. We can also define the probability density function

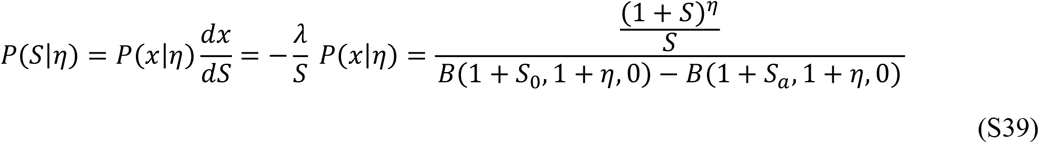

For the performance *Q*(*η*) we then use (S36-S39) to calculate the numerator of equation (S36)

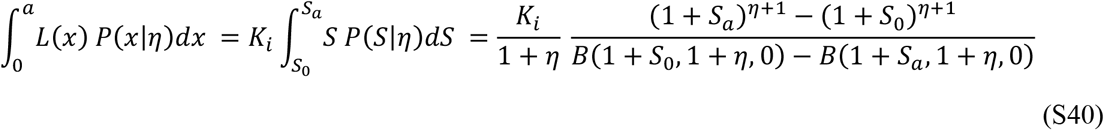

and the denominator

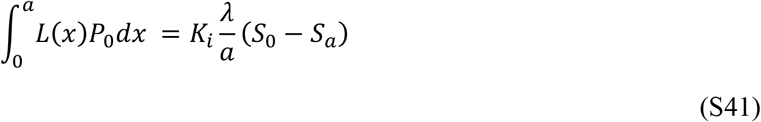

From this we obtain for the performance:

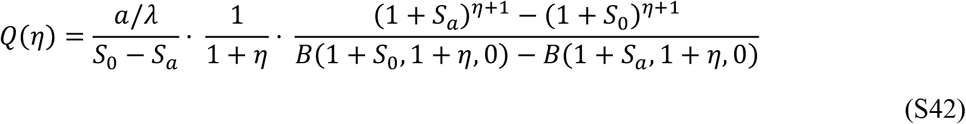

If we restrict the values of *η* to any interval, this performance has three regimes. At low values of *L*_*max*_ (or *S*_0_), this performance curve is convex (Figure 5B). At intermediate values, the curve is sigmoidal, and at high values, the curve is concave. Because of Jensen’s inequality, which states that for a convex function *Q*(*η*), that *Q*(⟨*η*⟩) ≤ ⟨*Q*(*η*)⟩, in the convex regime, it is possible for the average performance of a population with diverse *η* to exceed the performance of the average phenotype, ⟨*η*⟩. Conversely, in the concave regime, the population may underperform the average phenotype.

### QUANTIFICATION AND STATISTICAL ANALYSIS

All statistical testing was performed in Matlab (version R2020a). For each quantity, confidence intervals were calculated as described in the relevant figure caption. Data were quantified as described in the respective method subsections and/or figure captions.

## Supplemental Materials

**Figure S1.**
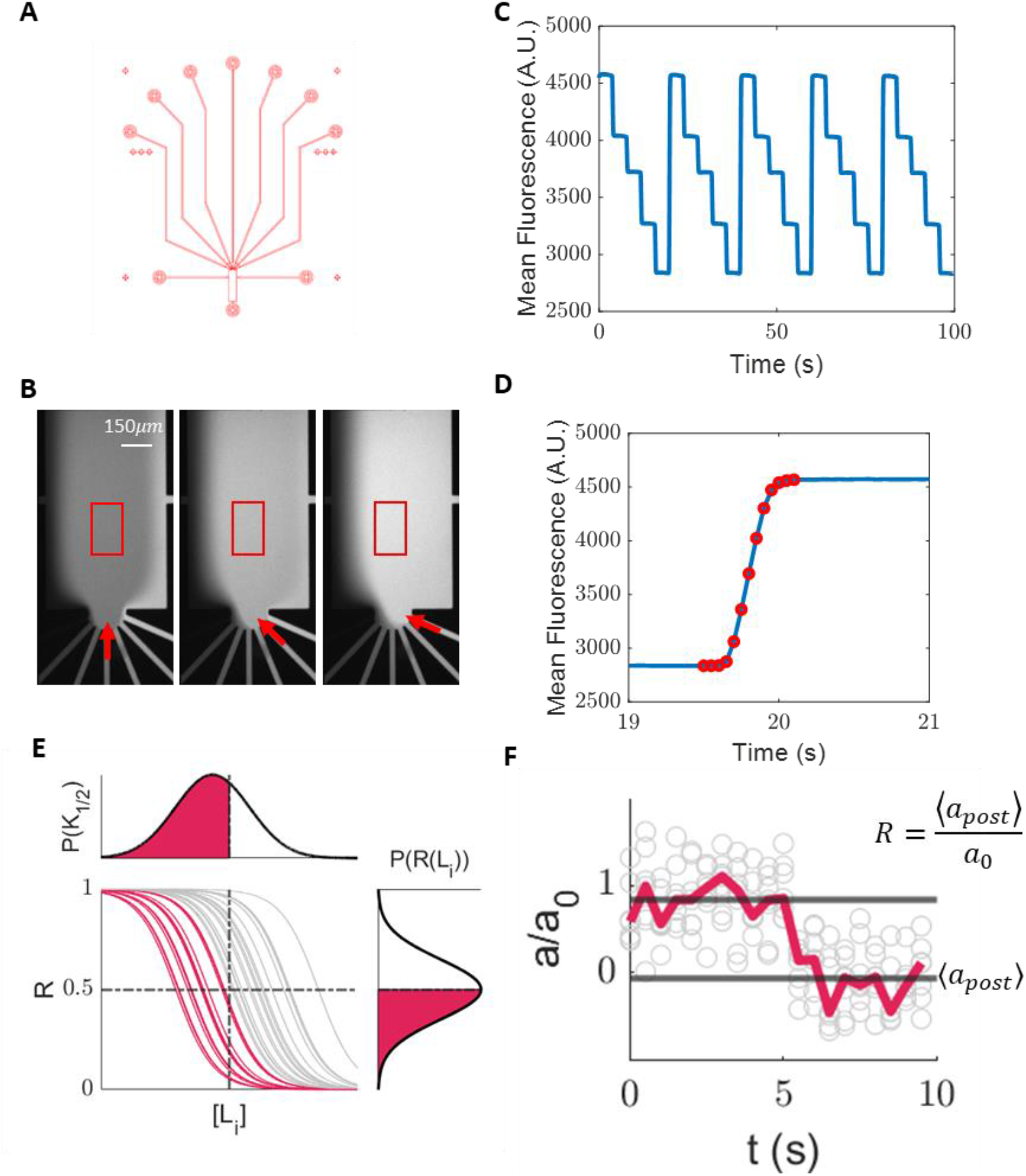
Seven-inlet microfluidic device developed to measure sensitivity distributions. **A)** Microfluidic device designed for single-cell FRET experiments. The device has seven inlets (top), two side outlets, and one main outlet (bottom). Before an experiment, cells are introduced through the main outlet, where they circle through the main chamber, and exit through one of the side outlets. The other side outlet is unused. **B)** Image of the main chamber with the approximate field of view boxed in red. Each inlet is loaded with different concentrations of fluorescein to visualize the flow. The stimulus concentration of the pressurized channel (red arrow) fully covers the field of view.**C)** Time series of fluorescence intensity measured at the field of view shown in B obtained by switching between channels filled with different fluoresceine concentrations over time. **D)** Higher resolution time series of fluorescence intensity upon switching pressurized channels. To fully turn over the solution in the field of view takes approximately 0.2 seconds. Each dot is the average fluorescence of the field of view for one frame, with images taken at 20 Hz. Interpolation of the points (blue line) is included as a visual guide. **E)** A schematic showing how the cumulative distribution function of the K_1/2_ can be calculated from single-cell responses to multiple stimulus levels. For a given stimulus level [L_i_], the response *R* is the post-stimulus kinase activity divided by the adapted kinase activity *a*_0_. The probability that a cell’s response is <0.5 equals the probability that the K_1/2_ is less than [L_i_]. **F)** An example single-cell average response is shown where grey dots are individual measurements of each response timepoint, while the red line is the median activity at each timepoint. Grey horizontal lines are the median kinase activity pre and post stimulus normalized to the steady-state activity. All seven measurements of the response are included to determine the average pre-stimulus activity, and average post-stimulus activity.

**Figure S2.**
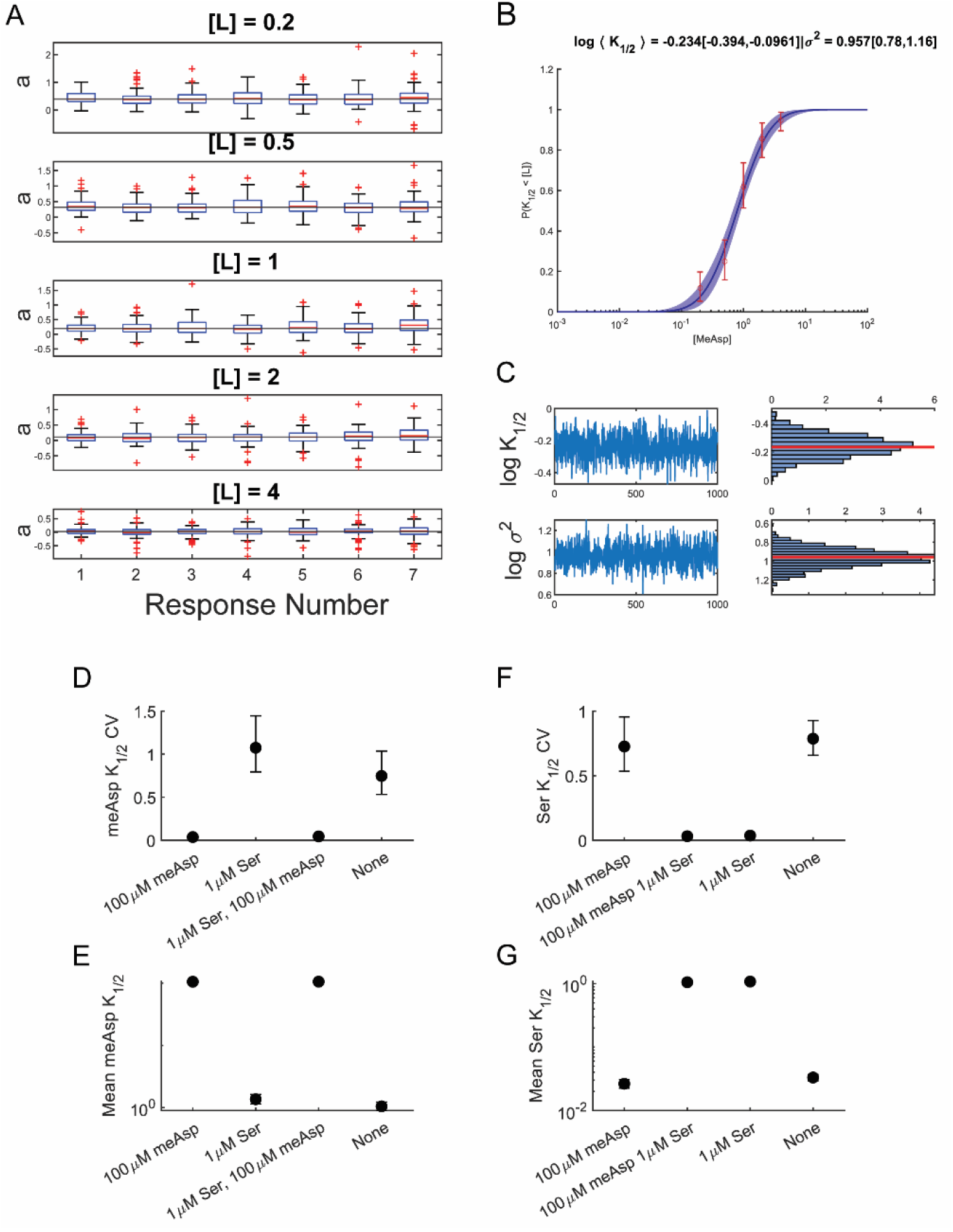
Statistical analysis of the *K*_1/2_ distribution. **(A)** Example distribution of post-stimulus kinase activity for seven responses to MeAsp at five stimulus concentrations. The response distribution is quasi-stationary over the course of the experiment, a necessary condition for our method to estimate the *K*_1/2_ distribution. **(B)** Parameters of the *K*_1/2_ cumulative distribution function (cdf) were determined by finding the maximum a posteriori parameter values of the log-normal cdf. Data points (open circles) are the fraction of response R<0.5 with error bars representing 95% confidence intervals (95 CI), calculated by bootstrapping over both the cells included in the analysis. The blue line is maximum a posteriori cdf with the 95 CI shaded in blue. **(C)** Uncertainty in the lognormal cdf parameters was determined by sampling the posterior with standard Markov-chain Monte Carlo methods with minimally informative priors. **(D-G)** Statistical analysis of *K*_1/2_ distributions in Figure 2. **(D)** CV and **(E)** mean *K*_1/2_ for MeAsp in four background conditions in Figure 2BC. **(F)** CV and **(G)** mean *K*_1/2_ for serine in the four background conditions in Figure 2DE. Error bars are 95-CI estimated using the posterior distribution of the fitted log-normal distribution parameters. Throughout the text, statistically significant differences occur only if there is no overlap in their 95% CI.

**Figure S3.**
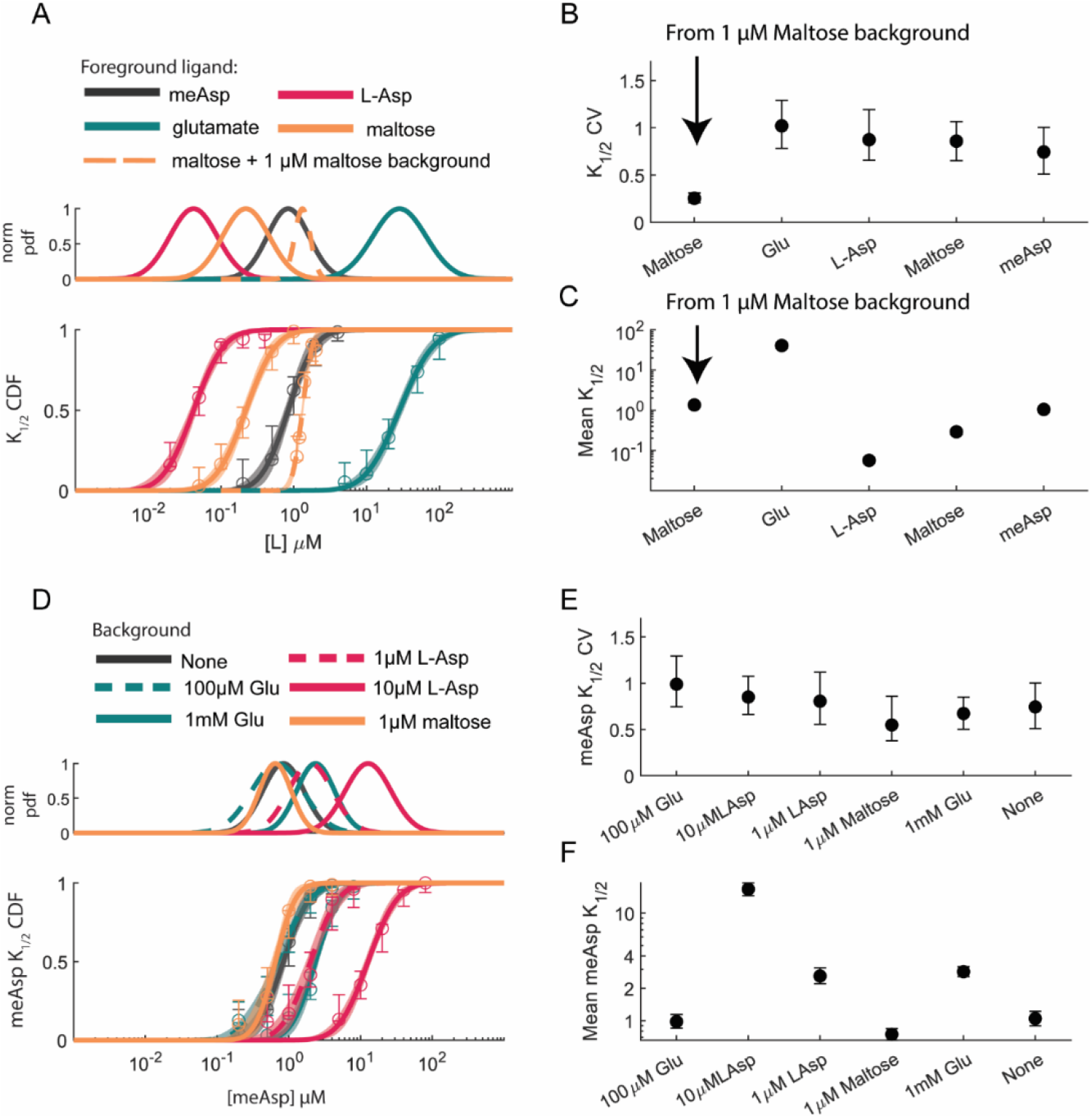
Statistical analysis of *K*_1/2_ distributions for Tar-binding ligands. **(A)** *K*_1/2_ distributions for four ligands that bind Tar in the absence of a background stimulus. The maltose *K*_1/2_ distribution is shown both with and without 1 µM maltose present. Statistical analysis of the mean **(B)** and CV **(C)** for each curve in panel A performed identically to Figure S2. Error bars are 95 CI. Only the CV for Maltose in 1 *µM* Maltose is significantly different from any other CV value. **(D)** Distribution of MeAsp *K*_1/2_ in the presence of various concentrations of glutamate, L-aspartate, and maltose. Statistical analysis of the **(E)** CV and **(F)** mean for the MeAsp *K*_1/2_ after adaptation to various competitive background stimuli. Error bars represent 95-CI. Presence of ligands that compete with MeAsp for binding sites do not affect the *K*_1/2_ CV, only the mean.

**Figure S4.**
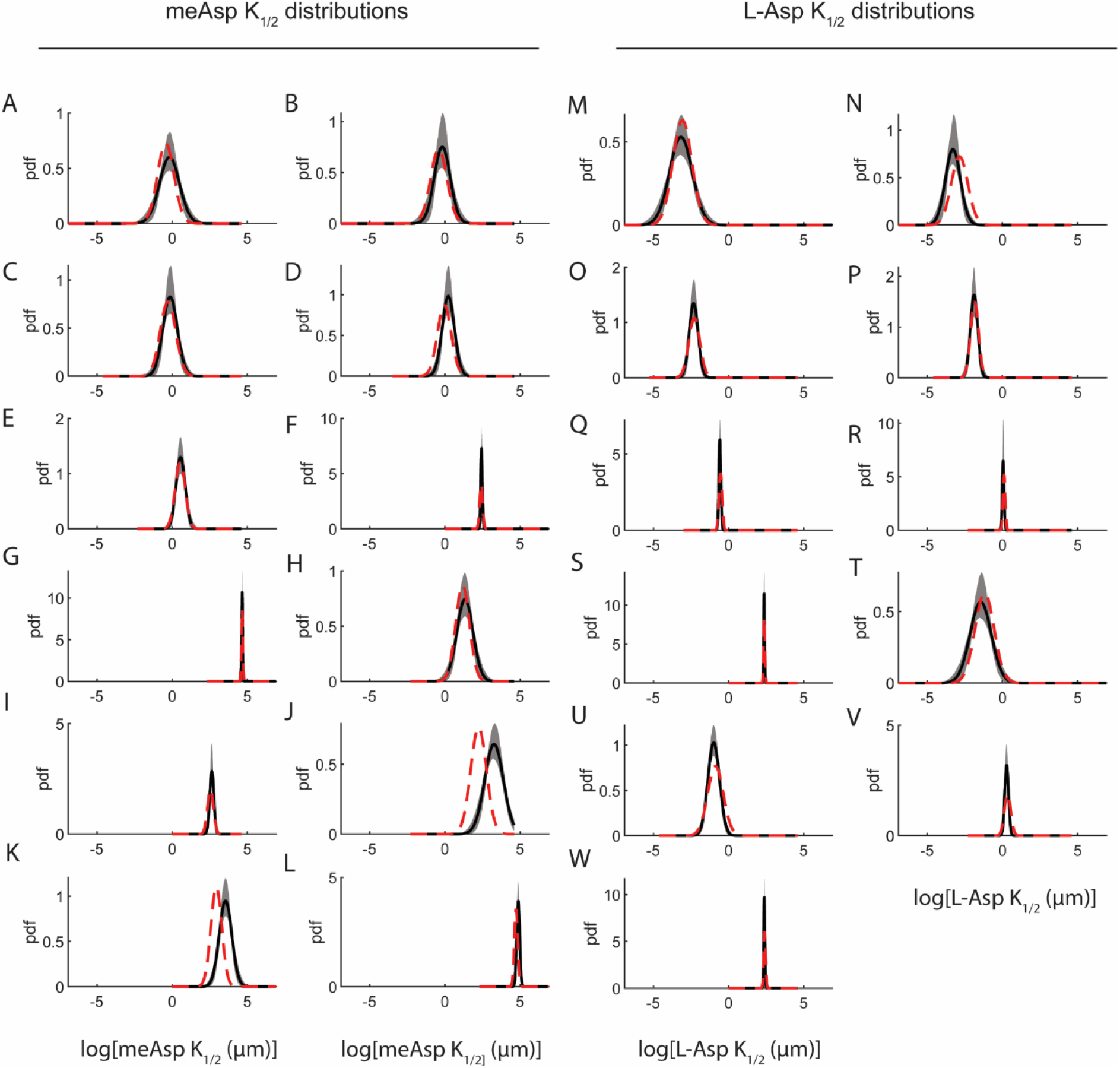
Comparison of measured and predicted *K*_1/2_ distributions for MeAsp and L-asp in mixed backgrounds. **A-L**) Experimental MeAsp *K*_1/2_ distributions in black with 95-CI shown as the shaded region. Red dotted lines are the predicted MeAsp *K*_1/2_ distributions from the fit in Figure 4CD. Background conditions in each panel are **A)** no background, **B)** 0.01 *µM* MeAsp, **C)** 0.1 *µM* MeAsp, **D)** 0.3 *µM* MeAsp, **E)** 1 *µM* MeAsp, **F)** 10 *µM* MeAsp, **G)** 100 *µM* MeAsp, **H)** 1 *µM* L-asp + 1 *µM* MeAsp, **I)** 1 *µM* L-asp + 10 *µM* MeAsp, **J)** 10 *µM* L-asp + 1 *µM* MeAsp, **K)** 10 *µM* L-asp + 10 *µM* MeAsp, **L)** 10 *µM* L-asp + 100 *µM* MeAsp. **M-W)** Experimental and predicted L-asp *K*_1/2_ distributions. Background conditions in each panel are **M)** No background, **N)** 0.01 *µM* L-asp, **O)** 0.05 *µM* L-asp, **P)** 0.1 *µM* L-asp, **Q)** 0.5 *µM* L-asp, **R)** 1 *µM* L-asp, **S)** 10 *µM* L-asp, **T)** 100 *µM* MeAsp, **U)** 100 *µM* MeAsp + 0.1 *µM* L-asp, **V)** 100 *µM* MeAsp + 1 *µM* L-asp, **W)** 100 *µM* MeAsp + 10 *µM* L-asp.

**Figure S5.**
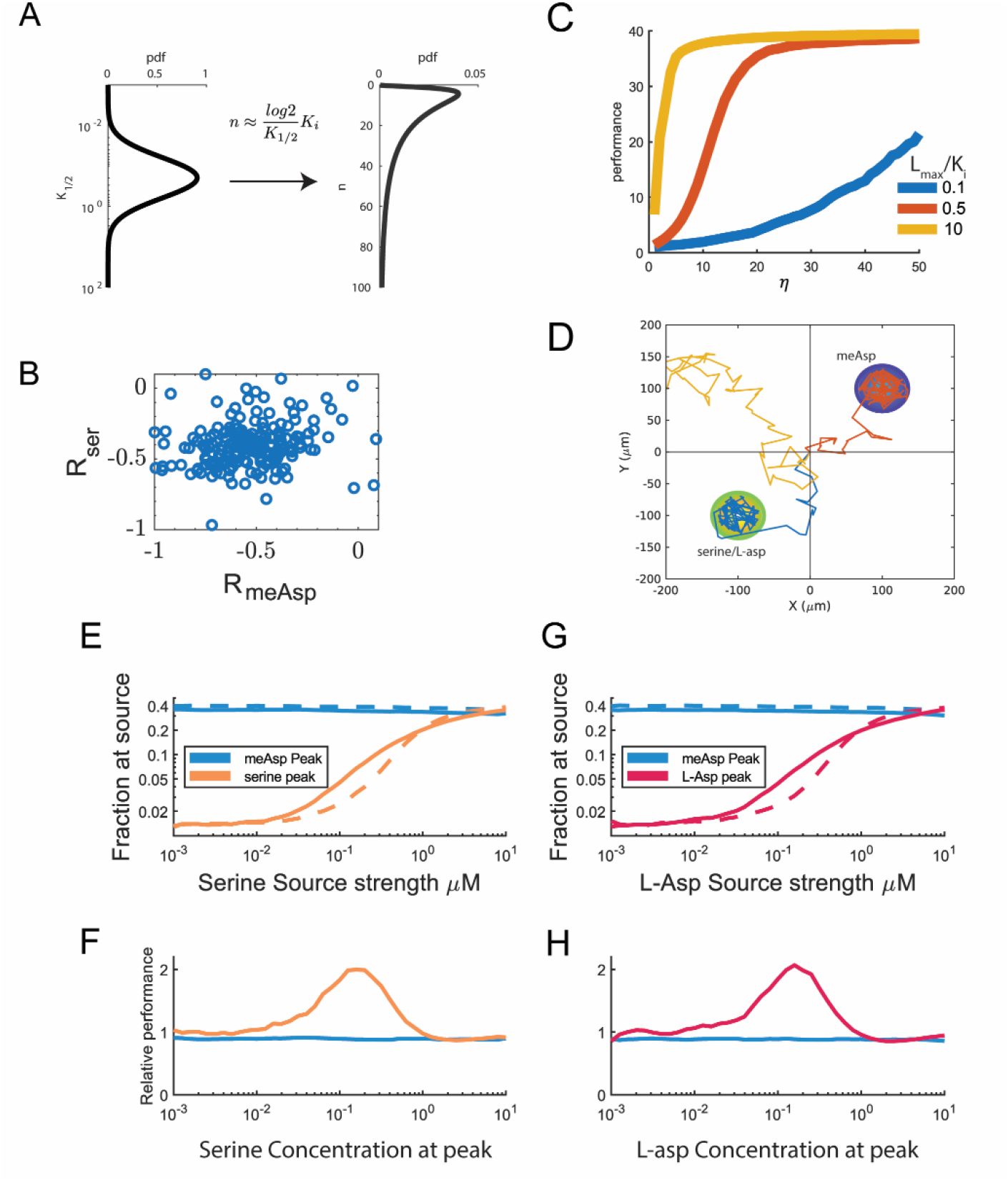
**A)** The distribution of receptor cooperativity *n* in the agent-based simulation can be extracted from the 0-background MeAsp *K*_1/2_ as consequence of the MWC model. **B)** Response amplitudes to 1uM MeAsp and 0.1 uM serine are uncorrelated, so in our simulations, when multiple receptors were considered, we allowed *n*_*Tar*_ and *n*_*Tsr*_ to be uncorrelated. **C)** Steady-state performance as a function of *η* = *nχ*/*D* in a single gaussian stimulus, determined numerically by MCMC sampling of the cell density equation. Performance is defined as average ligand concentration across the population density. **D)** Environment for agent-based simulation with two separate gaussian stimulus peaks. The MeAsp peak is fixed at 100*µM*. The second peak contains either serine or L-asp at variable peak concentration. Both gradients have the same length scale. Three example behavior trajectories are shown. **E)** Accumulation at a serine or MeAsp peak for a diverse population (solid line) and a population with uniform *n*_*Tar*_ and *n*_*Tsr*_ (dotted line), as described in main text. **F)** Relative performance of diverse population to the homogenous population at the serine and MeAsp peaks. The diverse population gains up to a 2-fold advantage for intermediate concentrations of serine. **G-H)** Same measurements as E-F but with the serine peak replaced with an L-asp peak with the same statistics. Even with this competitive attractant peak, the diverse population has up to a 2-fold advantage of the homogenous population at intermediate L-asp source strength.

**Table S1.** Optimal fitting parameters for the *K*_1/2_ CV model.

| Parameter | Value ± std deviation |
| --- | --- |
| $K_0$ (MeAsp) | $1.29 \pm 0.20 \mu\text{M}$ |
| $\sigma_0$ (MeAsp) | $0.84 \pm 0.37 \mu\text{M}$ |
| $K_i$ (MeAsp) | $18.1 \pm 0.98 \mu\text{M}$ |
| $K_0$ (L-aspartate) | $0.057 \pm 0.005 \mu\text{M}$ |
| $\sigma_0$ (L-aspartate) | $0.041 \pm 0.020 \mu\text{M}$ |
| $K_i$ (L-aspartate) | $0.86 \pm 0.10 \mu\text{M}$ |

## References

1. Qin, S., Li, Q., Tang, C., and Tu, Y. (2019). Optimal compressed sensing strategies for an array of nonlinear olfactory receptor neurons with and without spontaneous activity. Proceedings of the National Academy of Sciences 116, 20286–20295. 10.1073/pnas.1906571116.

2. Kadakia, N., and Emonet, T. (2019). Front-end Weber-Fechner gain control enhances the fidelity of combinatorial odor coding. eLife 8. 10.7554/elife.45293.

3. Pfister, P., Smith, B.C., Evans, B.J., Brann, J.H., Trimmer, C., Sheikh, M., Arroyave, R., Reddy, G., Jeong, H.-Y., Raps, D.A., et al. (2020). Odorant Receptor Inhibition Is Fundamental to Odor Encoding. Current Biology 30, 2574-2587.e2576. 10.1016/j.cub.2020.04.086.

4. Hallem, E.A., and Carlson, J.R. (2006). Coding of Odors by a Receptor Repertoire. Cell 125, 143–160. 10.1016/j.cell.2006.01.050.

5. Narla, A.V., Borenstein, D.B., and Wingreen, N.S. (2021). A biophysical limit for quorum sensing in biofilms. Proceedings of the National Academy of Sciences 118, e2022818118. 10.1073/pnas.2022818118.

6. Hurley, A., and Bassler, B.L. (2017). Asymmetric regulation of quorum-sensing receptors drives autoinducer-specific gene expression programs in Vibrio cholerae. PLOS Genetics 13, e1006826. 10.1371/journal.pgen.1006826.

7. Parkinson, J.S., Hazelbauer, G.L., and Falke, J.J. (2015). Signaling and sensory adaptation in Escherichia coli chemoreceptors: 2015 update. Trends in microbiology 23, 257–266. 10.1016/j.tim.2015.03.003.

8. Waite, A.J., Frankel, N.W., and Emonet, T. (2018). Behavioral Variability and Phenotypic Diversity in Bacterial Chemotaxis. Annual Review of Biophysics 47, 595–616. 10.1146/annurev-biophys-062215-010954.

9. Vaknin, A., and Berg, H.C. (2008). Direct evidence for coupling between bacterial chemoreceptors. J Mol Biol 382, 573–577. 10.1016/j.jmb.2008.07.026.

10. Piñas, G.E., Frank, V., Vaknin, A., and Parkinson, J.S. (2016). The source of high signal cooperativity in bacterial chemosensory arrays. Proceedings of the National Academy of Sciences 113, 3335–3340. 10.1073/pnas.1600216113.

11. Sourjik, V., and Berg, H.C. (2002). Receptor sensitivity in bacterial chemotaxis. Proc. Natl. Acad. Sci. USA 99, 123–127. 10.1073/pnas.011589998.

12. Sourjik, V., and Berg, H.C. (2004). Functional interactions between receptors in bacterial chemotaxis. Nature 428, 437–441. 10.1038/nature02406.

13. Meir, Y., Jakovljevic, V., Oleksiuk, O., Sourjik, V., and Wingreen, N.S. (2010). Precision and Kinetics of Adaptation in Bacterial Chemotaxis. Biophys. J. 99, 2766–2774. 10.1016/j.bpj.2010.08.051.

14. Sanders, D.A., and Koshland, D.E. (1988). Receptor interactions through phosphorylation and methylation pathways in bacterial chemotaxis. Proceedings of the National Academy of Sciences 85, 8425–8429. 10.1073/pnas.85.22.8425.

15. Hansen, C.H., Sourjik, V., and Wingreen, N.S. (2010). A dynamic-signaling-team model for chemotaxis receptors in Escherichia coli. Proc Natl Acad Sci U S A 107, 17170–17175. 10.1073/pnas.1005017107.

16. Lan, G., Schulmeister, S., Sourjik, V., and Tu, Y. (2011). Adapt locally and act globally: strategy to maintain high chemoreceptor sensitivity in complex environments. Mol Syst Biol 7, 475. 10.1038/msb.2011.8.

17. Karin, O., and Alon, U. (2021). Temporal fluctuations in chemotaxis gain implement a simulated-tempering strategy for efficient navigation in complex environments. iScience 24, 102796. 10.1016/j.isci.2021.102796.

18. Frankel, N.W., Pontius, W., Dufour, Y.S., Long, J., Hernandez-Nunez, L., and Emonet, T. (2014). Adaptability of non-genetic diversity in bacterial chemotaxis. Elife 3. 10.7554/eLife.03526.

19. Salek, M.M., Carrara, F., Fernandez, V., Guasto, J.S., and Stocker, R. (2019). Bacterial chemotaxis in a microfluidic T-maze reveals strong phenotypic heterogeneity in chemotactic sensitivity. Nature Communications 10. 10.1038/s41467-019-09521-2.

20. Colin, R., Rosazza, C., Vaknin, A., and Sourjik, V. (2017). Multiple sources of slow activity fluctuations in a bacterial chemosensory network. Elife 6. 10.7554/eLife.26796.

21. Keegstra, J.M., Kamino, K., Anquez, F., Lazova, M.D., Emonet, T., and Shimizu, T.S. (2017). Phenotypic diversity and temporal variability in a bacterial signaling network revealed by single-cell FRET. Elife 6. 10.7554/eLife.27455.

22. Mattingly, H.H., Kamino, K., Machta, B.B., and Emonet, T. (2021). Escherichia coli chemotaxis is information limited. Nat Phys. 10.1038/s41567-021-01380-3.

23. Kamino, K., Kadakia, N., Avgidis, F., Liu, Z.X., Aoki, K., Shimizu, T.S., and Emonet, T. (2023). Optimal inference of molecular interaction dynamics in FRET microscopy. Proc Natl Acad Sci U S A 120, e2211807120. 10.1073/pnas.2211807120.

24. Korobkova, E., Emonet, T., Vilar, J.M., Shimizu, T.S., and Cluzel, P. (2004). From molecular noise to behavioural variability in a single bacterium. Nature 428, 574–578. 10.1038/nature02404.

25. Park, H., Pontius, W., Guet, C.C., Marko, J.F., Emonet, T., and Cluzel, P. (2010). Interdependence of behavioural variability and response to small stimuli in bacteria. Nature 468, 819–823. 10.1038/nature09551.

26. Moore, J.P., Kamino, K., and Emonet, T. (2021). Non-Genetic Diversity in Chemosensing and Chemotactic Behavior. International Journal of Molecular Sciences 22, 6960. 10.3390/ijms22136960.

27. Kamino, K., Keegstra, J.M., Long, J., Emonet, T., and Shimizu, T.S. (2020). Adaptive tuning of cell sensory diversity without changes in gene expression. Science Advances 6, eabc1087. 10.1126/sciadv.abc1087.

28. Koler, M., Parkinson, J.S., and Vaknin, A. (2024). Signal integration in chemoreceptor complexes. Proceedings of the National Academy of Sciences 121. 10.1073/pnas.2312064121.

29. Neumann, S., Hansen, C.H., Wingreen, N.S., and Sourjik, V. (2010). Differences in signalling by directly and indirectly binding ligands in bacterial chemotaxis. The EMBO journal 29, 3484–3495. 10.1038/emboj.2010.224.

30. Yang, Y., M. Pollard A., Höfler, C., Poschet, G., Wirtz, M., Hell, R., and Sourjik, V. (2015). Relation between chemotaxis and consumption of amino acids in bacteria. Molecular Microbiology 96, 1272–1282. 10.1111/mmi.13006.

31. Frank, V., Piñas, G.E., Cohen, H., Parkinson, J.S., and Vaknin, A. (2016). Networked Chemoreceptors Benefit Bacterial Chemotaxis Performance. mBio 7, e01824–01816. 10.1128/mbio.01824-16.

32. Phillips, R. (2020). The molecular switch: signaling and allostery, 1. Edition (Princeton University Press).

33. Tu, Y. (2013). Quantitative modeling of bacterial chemotaxis: signal amplification and accurate adaptation. Annu Rev Biophys 42, 337–359. 10.1146/annurev-biophys-083012-130358.

34. Mello, B.A., and Tu, Y. (2005). An allosteric model for heterogeneous receptor complexes: Understanding bacterial chemotaxis responses to multiple stimuli. Proceedings of the National Academy of Sciences 102, 17354–17359. 10.1073/pnas.0506961102.

35. Keymer, J.E., Endres, R.G., Skoge, M., Meir, Y., and Wingreen, N.S. (2006). Chemosensing in Escherichia coli: Two regimes of two-state receptors. Proceedings of the National Academy of Sciences 103, 1786–1791. 10.1073/pnas.0507438103.

36. Pinas, G.E., DeSantis, M.D., Cassidy, C.K., and Parkinson, J.S. (2022). Hexameric rings of the scaffolding protein CheW enhance response sensitivity and cooperativity in Escherichia coli chemoreceptor arrays. Sci Signal 15, eabj1737. 10.1126/scisignal.abj1737.

37. Barkai, N., and Leibler, S. (1997). Robustness in simple biochemical networks. Nature 387, 913–917. 10.1038/43199.

38. Yi, T.M., Huang, Y., Simon, M.I., and Doyle, J. (2000). Robust perfect adaptation in bacterial chemotaxis through integral feedback control. P Natl Acad Sci USA 97, 4649–4653. DOI 10.1073/pnas.97.9.4649.

39. Somavanshi, R., Ghosh, B., and Sourjik, V. (2016). Sugar Influx Sensing by the Phosphotransferase System of Escherichia coli. PLOS Biology 14, e2000074. 10.1371/journal.pbio.2000074.

40. Shoval, O., Goentoro, L., Hart, Y., Mayo, A., Sontag, E., and Alon, U. (2010). Fold-change detection and scalar symmetry of sensory input fields. Proc Natl Acad Sci U S A 107, 15995–16000. 10.1073/pnas.1002352107.

41. Fu, X., Kato, S., Long, J., Mattingly, H.H., He, C., Vural, D.C., Zucker, S.W., and Emonet, T. (2018). Spatial self-organization resolves conflicts between individuality and collective migration. Nature Communications 9. 10.1038/s41467-018-04539-4.

42. Li, L., Zhang, X., Sun, Y., Ouyang, Q., Tu, Y., and Luo, C. (2024). Phenotypic Variability Shapes Bacterial Responses to Opposing Gradients. PRX Life 2. 10.1103/prxlife.2.013001.

43. Keller, E.F., and Segel, L.A. (1971). Traveling bands of chemotactic bacteria: a theoretical analysis. J Theor Biol 30, 235–248. 10.1016/0022-5193(71)90051-8.

44. Jensen, J.L.W.V. (1906). Sur les fonctions convexes et les inégalités entre les valeurs moyennes. Acta Mathematica 30, 175–193. 10.1007/bf02418571.

45. Long, J., Zucker, S.W., and Emonet, T. (2017). Feedback between motion and sensation provides nonlinear boost in run-and-tumble navigation. PLOS Computational Biology 13, e1005429. 10.1371/journal.pcbi.1005429.

46. Dufour, Y.S., Gillet, S., Frankel, N.W., Weibel, D.B., and Emonet, T. (2016). Direct Correlation between Motile Behavior and Protein Abundance in Single Cells. PLOS Computational Biology 12, e1005041. 10.1371/journal.pcbi.1005041.

47. Hart, Y., Mayo, A.E., Shoval, O., and Alon, U. (2013). Comparing Apples and Oranges: Fold-Change Detection of Multiple Simultaneous Inputs. PLoS One 8, e57455. 10.1371/journal.pone.0057455.

48. Adler, M., and Alon, U. (2018). Fold-change detection in biological systems. Current Opinion in Systems Biology 8, 81–89. 10.1016/j.coisb.2017.12.005.

49. Qin, D., Xia, Y., and Whitesides, G.M. (2010). Soft lithography for micro- and nanoscale patterning. Nature Protocols 5, 491–502. 10.1038/nprot.2009.234.

50. Zal, T., and Gascoigne, N.R. (2004). Photobleaching-corrected FRET efficiency imaging of live cells. Biophys J 86, 3923–3939. 10.1529/biophysj.103.022087.

51. Tu, Y., Shimizu, T.S., and Berg, H.C. (2008). Modeling the chemotactic response of Escherichia coli to time-varying stimuli. Proceedings of the National Academy of Sciences 105, 14855–14860. 10.1073/pnas.0807569105.

52. Shimizu, T.S., Tu, Y., and Berg, H.C. (2010). A modular gradient-sensing network for chemotaxis in Escherichia coli revealed by responses to time-varying stimuli. Mol Syst Biol 6, 382. 10.1038/msb.2010.37.

53. Sneddon, M.W., Pontius, W., and Emonet, T. (2012). Stochastic coordination of multiple actuators reduces latency and improves chemotactic response in bacteria. Proc Natl Acad Sci U S A 109, 805–810. 10.1073/pnas.1113706109.

54. Dufour, Y.S., Fu, X., Hernandez-Nunez, L., and Emonet, T. (2014). Limits of Feedback Control in Bacterial Chemotaxis. PLoS Computational Biology 10, e1003694. 10.1371/journal.pcbi.1003694.

55. Neumann, S., Vladimirov, N., Krembel, A.K., Wingreen, N.S., and Sourjik, V. (2014). Imprecision of adaptation in Escherichia coli chemotaxis. PLoS One 9, e84904. 10.1371/journal.pone.0084904.

56. Wong-Ng, J., Melbinger, A., Celani, A., and Vergassola, M. (2016). The Role of Adaptation in Bacterial Speed Races. PLoS Comput Biol 12, e1004974. 10.1371/journal.pcbi.1004974.

57. Tu, Y., and Rappel, W.J. (2018). Adaptation of Living Systems. Annu Rev Condens Matter Phys 9, 183–205. 10.1146/annurev-conmatphys-033117-054046.

58. Goentoro, L., Shoval, O., Kirschner, M.W., and Alon, U. (2009). The Incoherent Feedforward Loop Can Provide Fold-Change Detection in Gene Regulation. Mol Cell 36, 894–899. 10.1016/j.molcel.2009.11.018.

59. Tyson, J.J., Chen, K.C., and Novak, B. (2003). Sniffers, buzzers, toggles and blinkers: dynamics of regulatory and signaling pathways in the cell. Current Opinion in Cell Biology 15, 221–231. 10.1016/s0955-0674(03)00017-6.

